# PU.1 and MYC transcriptional network defines synergistic drug responses to KIT and LSD1 inhibition in acute myeloid leukemia

**DOI:** 10.1101/2021.08.24.456354

**Authors:** Brittany M. Smith, Jake VanCampen, Garth L. Kong, William Yashar, Yiu H. Tsang, Wesley Horton, Daniel J. Coleman, Joseph Estabrook, Theresa A. Lusardi, Gordon B. Mills, Brian J. Druker, Julia E. Maxson, Theodore P. Braun

## Abstract

Activating mutations in the KIT tyrosine receptor kinase confer an adverse prognosis for patients with acute myeloid leukemia (AML). Outside of bone marrow transplantation, treatment options are limited. Here we demonstrate combined KIT and LSD1 inhibition produces synergistic cell death against KIT mutant AML cells. This combination suppresses MYC expression to drive cell cycle exit and apoptosis. This decreased MYC expression results from a loss of PU.1 binding at downstream MYC enhancers. The drug combination also inactivates PI3K/AKT/GSK3a/b signaling to decrease MYC protein abundance. KIT-mutant AML cells rapidly adapt to KIT inhibitor monotherapy by restoring PI3K/AKT activity, but cannot when treated with combined KIT and LSD1 inhibitor. In addition, we validate MYC suppression as a mechanism of synergy in KIT-mutant AML patient samples. Collectively, this work provides rational for a clinical trial to assess the efficacy of KIT and LSD1 inhibition in patients with KIT-mutant AML.

**Statement of significance:** Effective treatment options for AML are limited. We describe the synergistic response to combined KIT and LSD1 inhibition in KIT-mutant AML and identify key biomarkers of drug response. The specificity and efficacy of this combination in cell lines and patient samples provides rationale for investigation in early phase clinical trials.

## Introduction

Acute myeloid leukemia (AML) results from the acquisition of two or more oncogenic mutations leading to proliferative advantage and hematopoietic differentiation block(1). Kinase mutations frequently provide this proliferative advantage, but are insufficient to produce AML in isolation(1). These mutations occur later in leukemogenesis, and are preceded by other genetic abnormalities(2–4). In contrast, mutations altering transcriptional or epigenetic regulatory proteins occur early in the disease course(4,5). The treatment of AML with kinase inhibitors has limited clinical efficacy due to short-lived responses and the inevitable development of drug resistance(6–8). Approaches to augment the efficacy of kinase inhibition by co-targeting of the leukemic epigenome represents a promising approach to obtain deeper and longer responses(9).

Epigenetic drugs have so far shown limited efficacy as monotherapy, with few patients demonstrating profound changes in disease volume(10–12). However, there is promising evidence showing increased efficacy when epigenetic drugs and kinase inhibitors are used in combination. Previously, we have showed that dual inhibition of JAK/STAT and an epigenetic regulator, lysine-specific demethylase 1 (LSD1) is an effective therapeutic strategy for CEBPA/CSF3R mutant AML(13). Inhibition of LSD1 restored differentiation-associated enhancer activity, further potentiating the action of JAK/STAT inhibitors in vitro and in vivo(13). Core binding factor (CBF) translocated AML shows a high degree of epigenetic similarity to CEBPA mutant AML resulting in a differentiation block, yet is associated with different signaling mutations(14,15). KIT mutations are the most common signaling mutation in CBF AML and drive proliferation via downstream JAK/STAT and MAPK pathway activation(16–18). CBF AML alone is favorable risk, however, the presence of KIT mutations is associated with an increased risk of relapse(16,19). Avapritinib (BLU-285) is a KIT inhibitor that is currently approved by the US Food and Drug Administration (FDA) for advanced systemic mastocytosis and gastrointestinal stromal tumor (GIST), both of which have activating KIT mutations (NCT00782067, NCT02508532). Given the short-lived responses to therapeutic inhibition of other kinases in AML, avapritinib monotherapy is unlikely to produce durable responses in KIT-mutant AML. However, given the efficacy of kinase plus LSD1 inhibition other AML subtypes, this approach warrants investigation for the treatment of KIT-mutant AML.

Here, we demonstrate that LSD1 inhibition potentiates the activity of the KIT inhibitor, avapritinib, in leukemia cell lines and primary patient samples. This synergistic cytotoxicity is driven by perturbation in the MYC and PU.1 transcription factor networks, resulting in decreased expression of proliferation associated genes. These mechanistic findings provide a basis for extending this dual therapeutic strategy into other molecularly defined subtypes of AML. Additionally, we identified key biomarkers to assess drug efficacy in KIT-mutant AML.

## Results

### Characterization of synergistic cytotoxicity following KIT and LSD1 inhibition

Combined kinase and LSD1 inhibition shows promise as a therapeutic strategy in certain molecularly defined subsets of AML(13). To investigate this combined therapy strategy in KIT-mutant AML, we performed a drug synergy analysis on KIT-mutant AML cell lines with KIT and LSD1 inhibitors. This revealed LSD1 inhibition, with either GSK-LSD1 or ORY-1001, had minimal effect on cell viability as a single agent, but markedly potentiated the efficacy of avapritinib (Fig. 1A-C, Supplementary Fig. S1A-B). We confirmed these results in EOL-1 cells which are also sensitive to avapritinib due to having a PDGFRA translocation(20) (Supplementary Fig. S1C). A major concern of combination therapy is toxicity to normal tissues including the hematopoietic system. To evaluate the toxicity of this combination, we performed a hematopoietic colony forming assay using normal CD34+ cells in the presence of the single agents or the combination. LSD1 inhibition reduced colony formation by 33%, consistent with the know role of LSD1 in supporting multi-lineage hematopoiesis(21) (Fig. 1D). However, avapritinib did not depress colony formation alone and did not cause a further decrease in colony formation when combined with LSD1 inhibition. Furthermore, the drug combination failed to substantially alter the growth of leukemia cell lines without KIT mutations, arguing that the drug combination is specific to KIT-mutant cells (Fig. 1E). To further characterize the response to combined inhibition of KIT and LSD1, we performed a flow cytometry-based apoptosis assay. After 48 hours of treatment, the combination group contained more PI+ (dead) and Annexin V+/PI- (early apoptosis) cells versus DMSO (Supplementary Fig. S1D-E). These results confirm that the drug combination produces greater cytotoxicity than either drug alone. In other molecular subtypes of AML, LSD1 inhibition promotes cell death through differentiation of immature AML blasts(13,22–26). Surface expression of the maturation associated marker, CD86, was increased by LSD1 inhibitor alone and to a similar degree when combined with avapritinib (Supplementary Fig. S1F-G). However, this effect was modest and was not accompanied by significant morphologic differentiation after 72 hours of treatment (Supplementary Fig. S1H), in contrast to the effects seen in MLL-rearranged AML(25). Overall, LSD1 inhibition potentiates the cytotoxic effect of avapritinib in KIT-mutant AML cell lines. This effect is specific to KIT-mutant cells as the drug combination has limited impact on healthy bone marrow or KIT wild type leukemia lines. Finally, although there is some immunophenotypic evidence of differentiation with LSD1 inhibition, this is not modified by co-treatment with avapritinib, and is relatively modest compared with results obtained in MLL-rearranged models.

**Figure 1.**
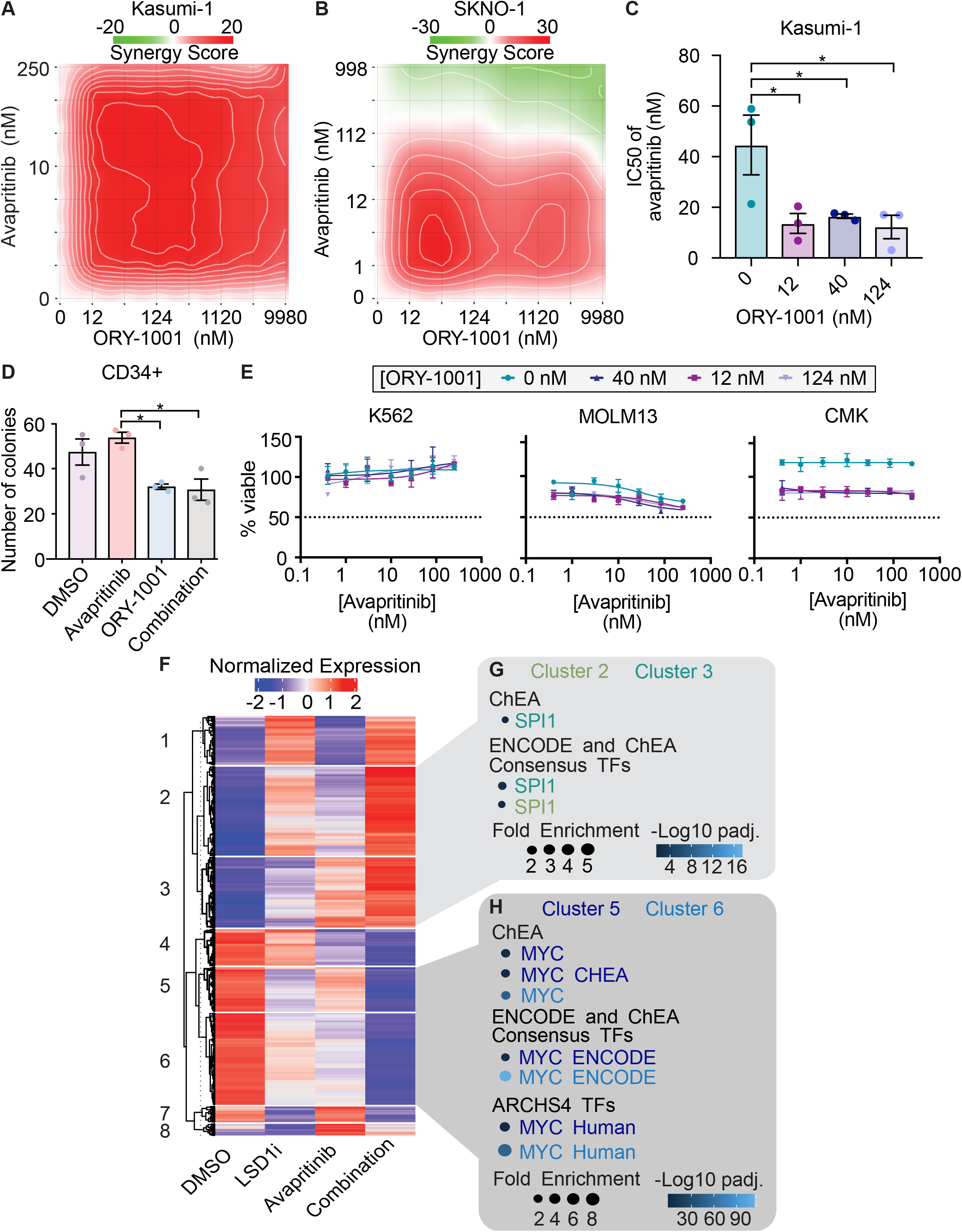
Synergistic cytotoxicity of dual KIT and LSD1 inhibition in a KIT-mutant AML cell line through activation of PU.1 and repression of MYC target genes. **A.** Drug matrix of Kasumi-1 cells treated for 72 h with avapritinib and ORY-1001 with synergy assessed by zero interaction potency (ZIP) score(74). **B.** Drug matrix of SKNO-1 cells treated for 72 h with avapritinib and ORY-1001 with synergy assessed by ZIP score. **C.** IC50 of avapritinib with different concentrations of ORY-1001 in Kasumi-1 cells treated for 72 h; one-way ANOVA with Holm-Sidak correction. **D.** Colony assay using healthy CD34+ cells in, treated for 14 days with avapritinib (12 nM) and/or ORY-1001 (12 nM); two-way ANOVA with Holm-Sidak correction. **E.** Viability assessment of K562, MOLM13, and CMK cells treated for 72 h with avapritinib and ORY-1001. **F.** Heatmap of differentially expressed genes from RNA-seq performed on Kasumi-1 cells treated with avapritinib (12 nM) and/or GSK-LSD1 (12 nM) for 12 h (*n*=3/group). Genes with significant differential expression are displayed by K-means clustering. **G, H.** Gene ontology analysis of clusters 2/3 and 5/6 from F, respectively. \**p* < 0.05

### Combined KIT and LSD1 inhibition leads to repression of MYC targets and activation of PU.1 targets

To investigate the mechanism of synergy between LSD1 and KIT inhibition, we performed RNA-sequencing (RNA-seq) on Kasumi-1 cells treated with avapritinib, GSK-LSD1, or the combination. Unsupervised clustering was performed on all differentially expressed genes to assess patterns of expression (Fig. 1F; Supplementary Tables S1-S2). Clusters 2 and 3 showed increased expression with the combination compared to either single agent, whereas clusters 5 and 6 showed decreased expression with the combination (Fig. 1G-H; Supplementary Table S3). We leveraged pathway analysis to identify biological programs that could drive the different expression patterns. Clusters 1 and 2 revealed enrichment for *SPI1* (PU.1) target genes, a transcription factor involved in hematopoietic differentiation (Fig. 1G)(27). Conversely, analysis of clusters 5 and 6 was highly enriched for MYC (a key regulator of proliferation and cell survival) target genes (Fig. 1H)(28). Taken together, increased expression of PU.1 target genes and downregulation of MYC target genes are features of the synergistic effect of avapritinib and LSD1 inhibition.

### Decreased activation of MYC bound promoters involved in cellular proliferation

Given the transcriptional impact of combined LSD1 and KIT inhibition on MYC target genes, we hypothesized MYC binding may be altered by the drug combination. To that end, we performed Cleavage Under Targets and Release Using Nuclease (CUT&RUN) to evaluate changes in MYC binding with drug treatment. While LSD1 inhibition alone modestly decreased global MYC binding, a greater decrease was observed with the drug combination (Fig. 2A). Consistent with this finding, MYC protein level also decreased with the drug combination (Supplementary Fig. S2A). The majority of high confidence MYC peaks localized to promoter regions, suggesting MYC-dependent transcriptional regulation commonly occurs at promoters (Fig. 2B). Given the decreased transcript abundance of MYC target genes (Fig. 1F), we hypothesized that MYC would predominantly localize to the transcription start sites (TSS) of down regulated genes. Indeed, MYC is enriched at the TSSs of genes with decreased expression in response to the drug combination (Fig. 2C). With LSD1 inhibition alone and combined KIT/LSD1 inhibition, MYC binding decreases at the TSSs of down regulated genes. To assess whether this loss of MYC binding was associated with decreased promoter activity, we assessed H3K27Ac read pileup at MYC bound promoters using cleavage under targets and tagmentation (CUT&Tag). Using H3K4me3 and H3K27Ac, we identified 9,369 active promoters (Supplementary Fig. S2B; Supplementary Table S4). With dual LSD1 and KIT inhibition there is a significant decrease in acetylation at MYC bound promoters (adj p-value < 2.2e-16) (Fig. 2D, Supplementary Fig. S2C; Supplementary Tables S5-S6). GO analysis of MYC bound promoters revealed an enrichment for proliferation related genes such as CDK13 and several ribosomal components (Fig. 2E, Supplementary Fig. S2D). Altogether, combined inhibition of LSD1 and KIT results in decreased total MYC protein, a loss of MYC binding, and decreased activation of genes involved in cell growth. To test if this altered gene expression program changes cell growth and division, we used a flow cytometry-based assay to evaluate changes in cell cycle. After 24, 48, and 72 hours of treatment, the drug combination significantly decreased the percentage of cycling cells and increased the percentage of cells in G1 phase (Supplementary Fig. S3A-B). Taken together, loss of MYC activity reduces cell cycle; however, it is unclear how LSD1 and KIT inhibition contribute to decreased MYC transcription.

**Figure 2.**
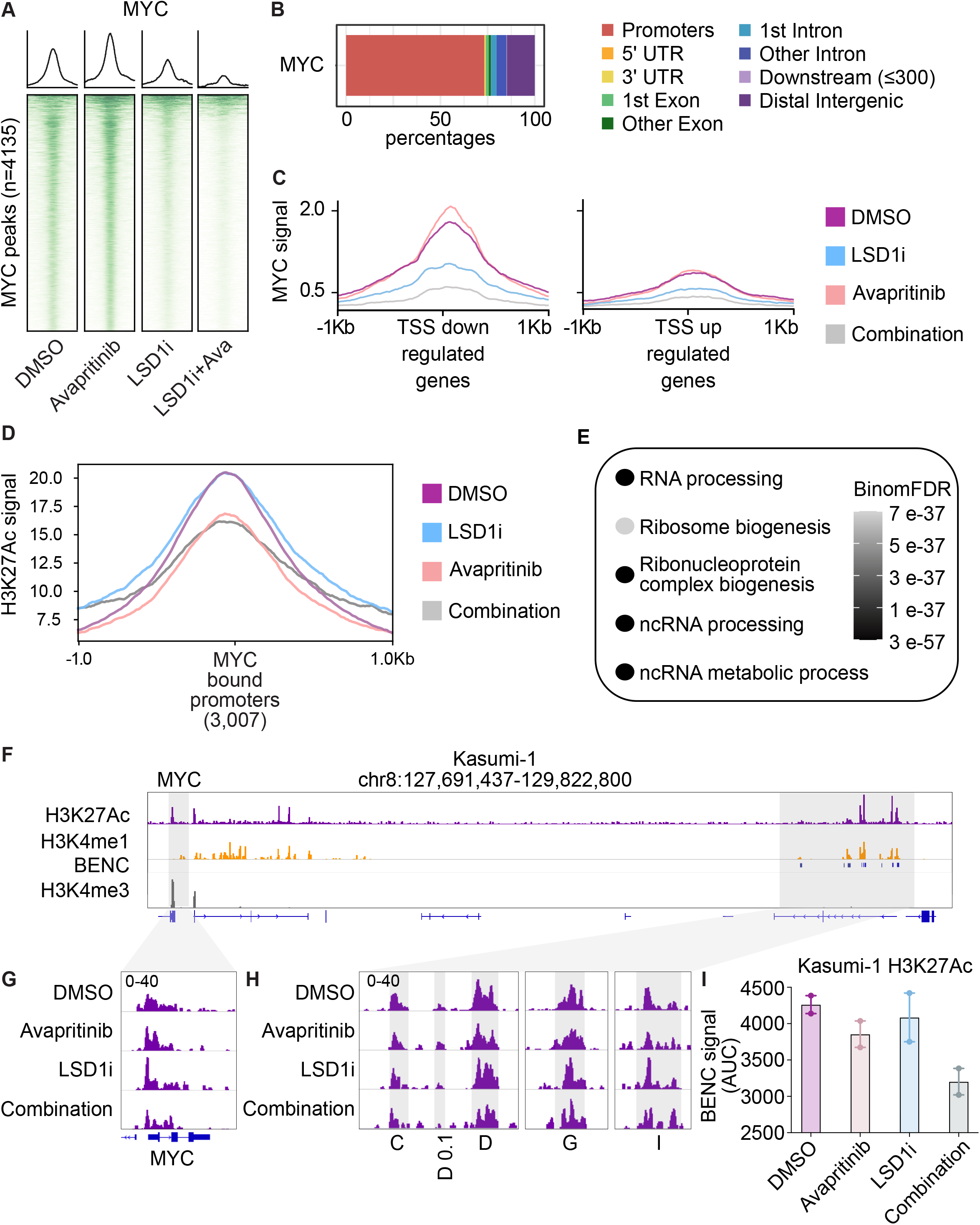
Repression of MYC bound promoters of cell cycle related programs. **A**. Kasumi-1 cells were treated for 24 h with avapritinib (12 nM) and/or GSK-LSD1 (12 nM; LSD1i) then subject to CUT&RUN (*n*=2/group). Heatmaps of global signal for MYC at high confidence consensus peaks (peak apex ± 1 kb). **B.** Annotation of consensus MYC peaks. **C.** MYC signal at TSSs of down or up regulated genes defined by RNA-seq. **D.** H3K27Ac signal at all MYC bound promoters in Kasumi-1 cells after 24 hours of treatment with avapritinib (12 nM) and/or GSK-LSD1 (12 nM; LSD1i). **E**. Gene ontology term enrichment for MYC bound promoters. **F.** Histone mark visualization with Integrative Genomics Viewer (IGV) at the MYC and blood enhancer cluster (BENC) locus. **G.** Histone acetylation in Kasumi-1 cells at the MYC locus. **H.** Histone acetylation at active BENC modules. **I.** AUC of acetylation signal at active BENC modules.

To gain further insight into the transcriptional control of MYC, we considered its promoter and enhancer regions. It has previously been shown in other AML models that MYC transcription is regulated by the blood enhancer cluster (BENC)(29,30). We used H3K4me1 signal and the previously annotated modules to identify the BENC in Kasumi-1 cells (Fig. 2F). H3K27Ac signal was used to identify active BENC modules (Fig. 2F). Indeed, acetylation was lost at the MYC promoter in Kasumi-1 cells following dual LSD1 and KIT inhibition (Fig. 2G). Acetylation signal within active BENC modules also decreased post drug treatment (Fig. 2H-I). Decreased BENC activity may contribute to the loss of MYC expression following KIT and LSD1 inhibition. In sum, LSD1 and KIT inhibition results in loss of MYC binding and decreased acetylation of proliferation associate promoters ultimately leading to cell cycle arrest.

### Combined KIT and LSD1 inhibition cause activation of GFI1 and LSD1 bound enhancers

The transcriptional response to combined KIT and LSD1 inhibition involves both repression of MYC target genes as well as activation of differentiation-associated genes. In other subtypes of AML, differentiation of AML blasts is a major driver of drug responses. A key driver of this differentiation response is displacement of the repressive transcription factor GFI1 from chromatin. GFI1 and LSD1 co-bind and pharmacologic inhibition of LSD1 results in a loss of GFI1 binding and activation of repressed enhancers of differentiation associated genes(23). To evaluate the role of GFI1 in the response to combined KIT and LSD1 inhibition in KIT-mutant AML, we assessed genome-wide binding of GFI1 and LSD1 by CUT&RUN. LSD1 binding did not globally change after treatment in any treatment condition (Supplementary Fig. S4A). GFI1 binding decreased with LSD1 inhibition (Supplementary Fig. S4B), but was lost to a greater degree with the drug combination. To assess whether this loss of GFI1 binding was associated with activation of the underlying enhancers, we used CUT&Tag to identify enhancers, marked by the presence of H3K4me1. Using this approach, we identified 4,199 active enhancers (Supplementary Fig. S2B; Supplementary Table 2). The enhancers bound by GFI1 and LSD1 showed subtle changes in the underlying H3K27Ac, however these changes were modest compared with the results seen in MLL rearranged AML treated with LSD1 inhibitors, arguing that other factors are the major drivers of the response to combined KIT and LSD1 (Supplementary Fig. S4C; Supplementary Tables S7-S8)(23). Indeed, this is consistent with the relatively modest immunophenotypic differentiation and absence of morphologic differentiation driven by drug treatment (Supplementary Fig. S1H).

### *LSD1 inhibition leads to loss of PU.1 at MYC* enhancers

PU.1 has also been implicated as a regulator of LSD1 inhibitor responses and PU.1 target genes are featured prominently in our RNA-seq analysis(25) (Fig. 1F). We therefore performed CUT&RUN to profile the genome wide binding of PU.1 (Fig. 3A). PU.1 binding is completely lost after treatment with LSD1 inhibition without changes in total PU.1 protein (Fig. 3A, Supplemental Fig. S5A). Annotation of PU.1 binding sites revealed localization outside of promoter regions (Supplemental Fig. S5B). To assess whether loss of PU.1 activity was sufficient to augment the cytotoxicity of KIT inhibition, we used both a pharmacologic inhibitor of PU.1 activity (DB2313) and a doxycycline-inducible short hairpin RNA (shRNA) targeting PU.1. In both instances, we found a dose-dependent increase in KIT inhibitor potency with increasing PU.1 inhibition or knockdown (KD) (Fig. 3B, Supplementary Fig. S5C-E). Next, we tested whether PU.1 KD and KIT inhibition recapitulates the transcriptional consequences of LSD1 and KIT inhibition. Bulk RNA-seq of Kasumi-1 cells treated with avapritinib and PU.1 shRNA showed a depletion of MYC target genes by gene set enrichment analysis (Fig. 3C; Supplementary Tables S9-S11). The genes from this MYC target geneset showed more repression with the combination than with either drug alone (Supplemental Fig. S5F). These data demonstrate that a loss of PU.1 activity contributes to the repression of MYC target genes.

**Figure 3.**
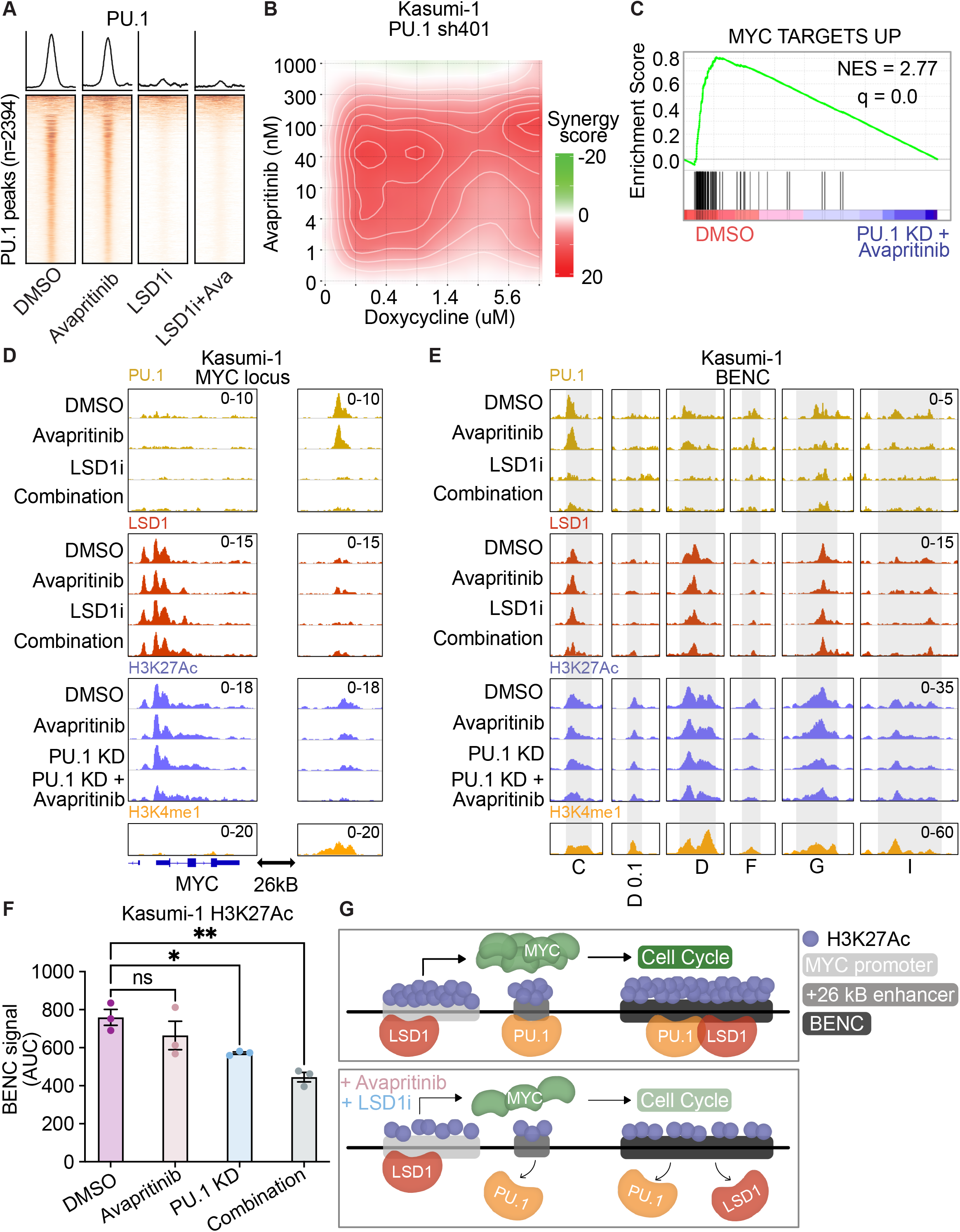
Loss of PU.1 binding at MYC enhancer resulting in loss of MYC enhancer and promoter activation. **A.** Kasumi cells treated for 24 h with avapritinib (12 nM) and/or GSK-LSD1 (12 nM; LSD1i) then subject to CUT&RUN for PU.1 (*n*=2/group). Heatmaps of global signal for PU.1 at high confidence consensus PU.1 peaks (peak apex ± 1 kb). **B**. Drug matrix of avapritinib and doxycycline on Kasumi-1 PU.1 sh401 cells treated for 72 h. Synergy assessed by ZIP scores. **C.** Depleted gene sets from bulk RNA-seq on PU.1 sh401, induced with doxycycline (1 μg/mL) 48 h before treatment with avapritinib (50 nM) for 24 h. NES = normalized enrichment score (*q* < 0.05). GSEA *p* value calculated by empirical permutation test and FDR adjusted. **D, E**. Kasumi-1 cells treated for 24 h with avapritinib (50 nM) and/or doxycycline (1 μg/mL) to induce PU.1 knockdown were used to perform CUT&Tag for H3K27Ac (*n*=3/group). H3K4me1, PU.1, and LSD1 signal from above datasets in Kasumi-1 cells. Visualization of +26 Kb MYC and blood super-enhancer cluster (BENC). **F**. Quantification of cumulative AUC of H3K27Ac signal at active BENC modules; one-way ANOVA with Holms-Sidak correction. **G.** Model describing loss of PU.1 binding after dual LSD1 and KIT inhibition at MYC +26 kB enhancer and BENC. PU.1 no longer activates MYC promoter resulting in decreased MYC protein, leading to decreased expression of MYC target genes including those involved with cell proliferation. * *p* < 0.05, \*\**p* < 0.01

To understand how PU.1 KD regulates MYC expression, we performed CUT&Tag for H3K27Ac to look at activation of MYC regulatory elements. Acetylation of the MYC promoter significantly decreased with PU.1 KD and KIT inhibition (adj p-value = 0.03) (Fig. 3D; Supplementary Tables S12-S14). Interestingly, no PU.1 peak was identified at the MYC promoter. However, a PU.1 peak was identified at the annotated +26 Kb enhancer downstream of the MYC gene (31), which was marked by both H3K27Ac and H3K4me1 (Fig. 3D). Consistent with the global changes in PU.1 binding, PU.1 signal was lost with LSD1 inhibition at the +26 Kb MYC enhancer and PU.1 KD resulted in a significant loss of H3K27Ac which further decreased with KIT inhibition (adj p-value = 0.01; Fig. 3D). Interestingly, LSD1 and GFI1 are found at the MYC promoter, but they were not identified at the +26 Kb enhancer (Fig. 3D, Supplementary Fig. S5G). Moreover, LSD1 and PU.1 are both bound to active modules in the BENC, which loses acetylation with KIT inhibition and PU.1 KD (Fig. 3E-F). These data demonstrate that PU.1 may play a role in MYC transcription through downstream enhancers. When PU.1 binding is lost with LSD1 inhibition, BENC activation is reduced resulting in decreased transcription of MYC and its associated programs (Fig. 3G).

### KIT and LSD1 inhibition decrease MYC protein levels and repress LSD1 target genes through PI3K/AKT pathway

MYC is a key regulator of cellular proliferation and survival, among other functions, thus its transcription, activation, and stability are carefully regulated(32). KIT activates both MAPK/RAS/ERK and PI3K/AKT pathways each contributing to MYC protein abundance(33). We hypothesized KIT inhibition may alter MYC protein levels, leading to reduced MYC binding. To assess the dynamics of AKT and ERK pathway signaling after combined KIT and LSD1 inhibition, we used a novel fluorescent biosensor that enables live-cell imaging of AKT and ERK pathway activity via nuclear-cytoplasmic redistribution of fluorescent target proteins (Fig. 4A). We observed minimal change in ERK activity after either drug treatment, but observed notable changes in AKT activity (Fig. 4B, Supplementary Fig. S6A-C). To assess the relative contribution of each drug, we clustered cells based on their AKT dynamics after treatment (Fig. 4B, Supplementary Fig. S6D). KIT inhibition alone resulted in an increased number of cells demonstrating a steep drop in AKT activity with a gradual recovery to baseline over 24 hours (Cluster 5). In contrast, LSD1 inhibition increased the abundance of cells with gradually decreasing AKT activity during the observation period (Cluster 4). Combined inhibition of KIT and LSD1 resulted in the accumulation of cells demonstrating a sharp drop in AKT activity with minimal recovery (Cluster 6). These results collectively demonstrate that KIT inhibition results in a rapid decrease in AKT activity followed by subsequent slow return to baseline. Simultaneous inhibition of LSD1 attenuates this recovery with slow dynamics, consistent with a transcriptional response to drug.

**Figure 4.**
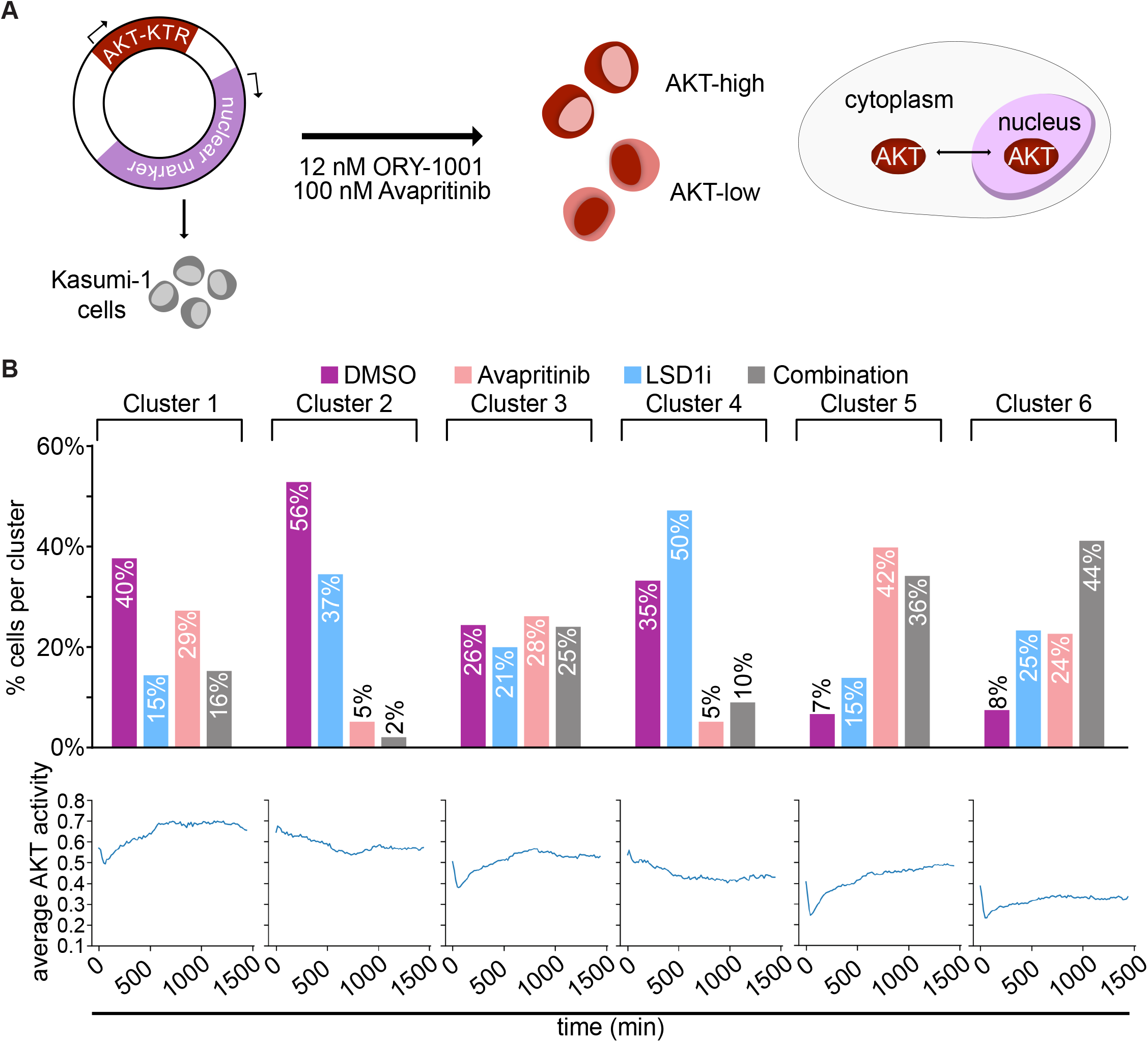
KIT and LSD1 inhibition attenuate AKT signaling. **A.** Experimental strategy to continuously assess AKT activity over time with AKT-KTR-mScarlet fluorescent biosensor. ERK activity was evaluated in the same way with ERK-KTR-Clover. **B.** Kasumi-1 cells were treated with avapritinib (100 nM) and/or ORY-1001 (12 nM). Subcellular localization of the fluorescent biosensors was captured with live-cell imaging. Cells were clustered based on AKT activity over time. Bar graph displays percentage of cells from each treatment group within each cluster.

To assess the changes in these signaling pathways at the phosphoproteomic level, we performed reverse phase protein array (RPPA) on Kasumi-1 cells treated with the single drugs or the combination. CausalPath phosphoprotein activity network revealed decreased activation of AKT upstream of decreased total MYC protein (Supplementary Fig. S7A-B; Supplementary Table S15). To understand the dynamics of AKT signaling after combined KIT and LSD1 inhibition, we used a heatmap to visualize individual members of the PI3K/AKT pathway(34–36) (Fig. 5 A-B; Supplementary Fig. S7C). Serine 21 of GSK3a/b is phosphorylated by AKT, resulting in inhibition of kinase activity (Fig. 5B). Active (dephosphorylated) GSK3a/b phosphorylates MYC to decrease protein abundance(37). After 1 h of treatment with KIT inhibition and the combination, total MYC protein and serine phosphorylated GSK3a/b (pS21) decreased markedly. Following KIT and LSD1 inhibition, PI3K/AKT is downregulated resulting in the reactivation of GSK3a/b, leading to decreased MYC protein.

**Figure 5.**
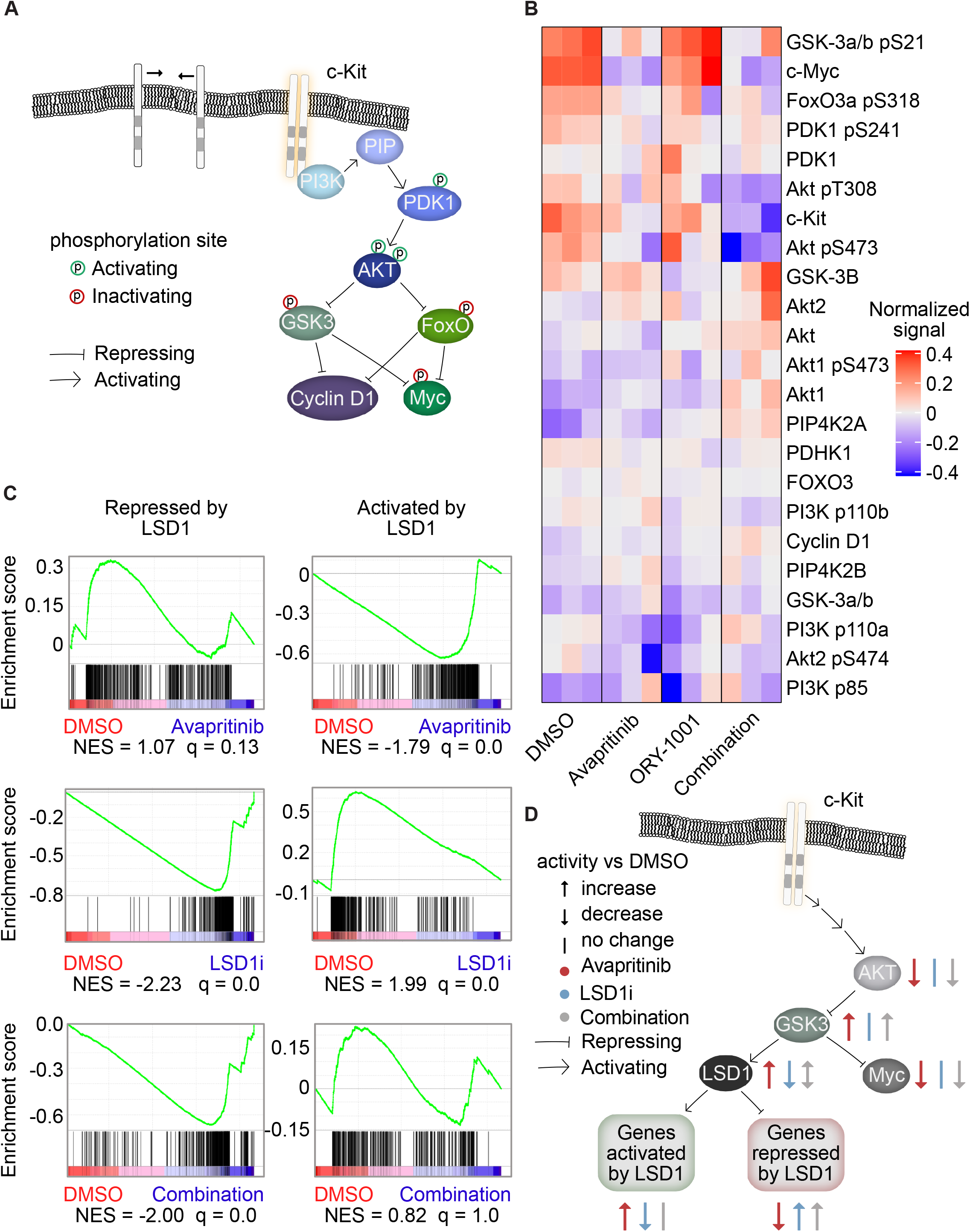
Coordinated PI3K/AKT signaling response to dual KIT and LSD1 inhibition. **A**. Depiction of PI3K/AKT signaling pathway(34–36). **B.** RPPA of Kasumi-1 cells treated for 1 h with avapritinib (100 nM) and/or ORY-1001 (12 nM). Heatmap of the normalized signal from PI3K/AKT pathway members. **C.** GSEA using genesets curated by bulk RNA-seq of Kasumi-1 cells treated for 12 h with GSK-LSD1 (12 nM). Genes repressed by LSD1 have significantly increased expression after LSD1 inhibition. Genes activated by LSD1 have significantly decreased expression after LSD1 inhibition. NES = normalized enrichment score (*q* < 0.05). GSEA *p* value calculated by empirical permutation test and FDR adjusted. **D.** Inhibition of KIT or LSD1 lead to opposing effect on LSD1 activity via the PI3K/AKT pathway. Dual inhibition results in repression of MYC and activation of genes natively repressed by LSD1.

As LSD1 inhibition potentiates the cytotoxic effects of avapritinib, we hypothesized that LSD1 might also interact with the KIT/AKT/GSK3a/b axis. Indeed, in glioblastoma cells (GSC11), GSK3b phosphorylates LSD1 resulting in activation(38). This finding suggests KIT inhibition could increase the activity of LSD1, potentially increasing the dependency on this pathway and rendering cells more sensitive to LSD1 inhibition. To test this, we identified two gene-sets of LSD1 responsive genes: genes that increase in expression with LSD1 inhibition (LSD1-repressed) and genes that decrease in expression with LSD1 inhibition (LSD1-activated). We then assessed gene set enrichment of these LSD1 responsive genes in cells responding to KIT inhibition. Consistent with the hypothesis that KIT and LSD1 inhibition exert opposing effects on LSD1 target genes, we observed that avapritinib resulted in activation of LSD1-activated genes and repression of LSD1-repressed genes (Fig. 5C; Supplementary Table S16). We next compared the effect of LSD1 inhibition alone to combined KIT and LSD1 inhibition. With or without KIT inhibition, LSD1 inhibition resulted in activation of LSD1-repressed genes. However, LSD1-activated genes were only repressed by LSD1-inhibitor monotherapy but not when combined with KIT inhibition. These data support a model in which KIT inhibition suppresses MYC activity but activates LSD1 as a drug escape mechanism. This adaptive response to KIT inhibition can be targeted with the addition of LSD1 inhibitor, potentiating cell death (Fig. 5D).

We also investigated signaling responses to KIT and LSD1 inhibition at a later timepoint. At 24h, we observed that avapritinib-treated cells demonstrated increased levels of phospho-AKT (both pS473 and pT308) (Supplementary Fig. S7C). Kit protein abundance was also upregulated at this time-point suggesting that upregulation of KIT may drive this later adaptive response. The addition of LSD1 inhibitor resulted in marked suppression of KIT protein abundance as well as phospho-AKT. Collectively, these results demonstrate a coordinated interplay between signaling pathways downstream of KIT (namely PI3K/AKT) and LSD1.

### LSD1 and KIT inhibition is synergistically cytotoxic in KIT-mutant patient samples through MYC network repression

To verify our findings in cell lines in primary AML samples, we identified thirteen samples from the Beat AML cohort with KIT mutations(39). Ten of the thirteen patient samples had mutations at D816, leading to ligand independent activation that is amenable to inhibition by avapritinib. Six of those patient samples had frozen viable cells available, four of which thawed with sufficient viability for further assays (Supplementary Table S17). We elected to evaluate transcriptional and epigenetic changes at 24 hours and drug synergy after 72 hours (Fig. 6A). Of the four patient samples, three responded with synergistic cytotoxicity to the LSD1 and KIT inhibitor (Fig. 6B, Supplementary Fig. S8A-C). Sample 17-00007 was from a relapsed patient and still showed synergistic response to LSD1 and KIT inhibition (Supplementary Fig. S8B). Sample 17-01023 did not respond, but had a KIT-mutant VAF of 6% arguing that this mutation only exists in a small subclone (Supplementary Fig. S8C). In sample 14-00613, bulk RNA-seq revealed downregulation of MYC target genes and cell cycle related genes (Fig. 6C-D; Supplementary Tables S18-S20). These transcriptional changes after dual drug treatment provide supporting evidence that suppression of MYC activity plays a key role in the response to LSD1 and KIT inhibition.

**Figure 6.**
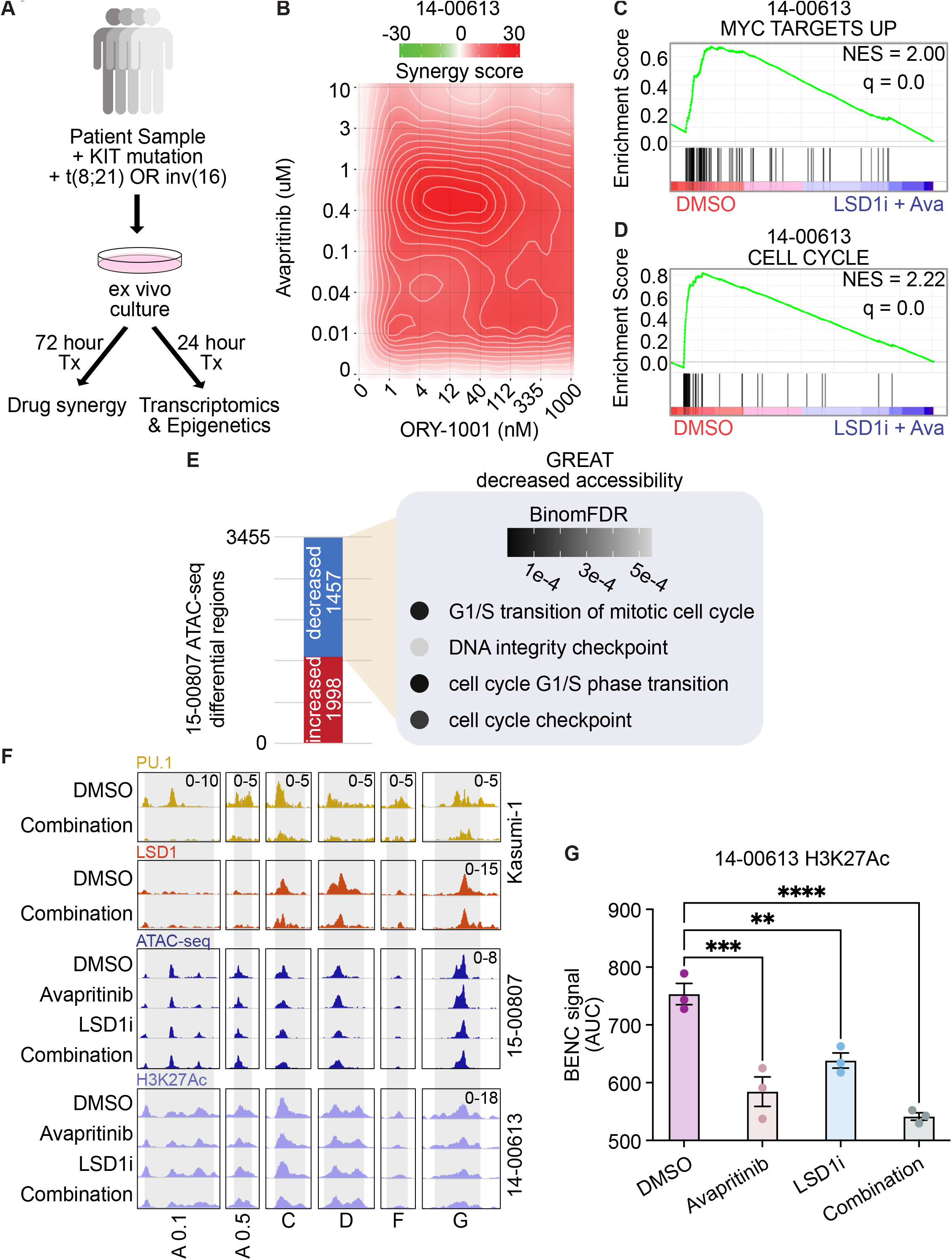
KIT and LSD1 inhibition synergistically target KIT-mutant AML patient samples resulting in decreased MYC and cell cycle programs. **A.** Experimental strategy for KIT-mutant patient samples. Frozen viable samples were cultured ex vivo and treated for 24 h before bulk RNA-seq and ATAC-seq. For synergy analysis, samples were drug treated for 72 hours before assessing drug synergy. **B.** Drug matrix of patient sample 14-00613 treated for 72 h with avapritinib and ORY-1001 with synergy assessed by ZIP score. **C, D.** Select depleted gene sets from bulk RNA-seq on 14-00613 treated with avapritinib (50 nM) and ORY-1001 (12 nM) or DMSO for 24 h. NES = normalized enrichment score (*q* < 0.05). GSEA *p* value calculated by empirical permutation test and FDR adjusted. **E.** Differential analysis of bulk ATAC-seq on 15-00807 treated with avapritinib (50 nM) and ORY-1001 (50 nM) compared to DMSO. Enrichment of GO terms for regions with significantly decreased accessibility. **F.** Visualization of Kasumi-1 PU.1 and LSD1 from above datasets, 15-00807 bulk ATAC-seq, and 14-00613 H3K27Ac at active BENC modules. H3K27Ac CUT&Tag was performed on 14-00613 following 24 hr treatment with avapritinib (350 nM) and/or ORY-1001 (12 nM). **G.** Quantification of 14-00613 H3K27Ac signal at active BENC modules by comparing AUC; one-way ANOVA with Holm-Sidak correction. \*\**p* < 0.01, *** *p* < 0.001, \*\*\*\**p* < 0.0001

Due to the epigenetic modifying role of LSD1, we were interested to know if dual inhibition of LSD1 and KIT altered the epigenetic landscape in patient samples. We used Transposase Accessible Chromatin using sequencing (ATAC-seq) to look at changes in chromatin accessibility in sample 15-00807(40). Differential accessibility analysis revealed 1,457 regions with significantly decreased accessibility after dual LSD1 and KIT inhibition (Fig. 6E). Gene Ontology analysis of these regions revealed enrichment for genes controlling cell cycle, including EEF2K, a kinase involved in cell cycle(41) (adj p-value = 0.0004) (Fig. 6E, Supplementary Fig. S8D; Supplementary Tables S21-S23). In order to specifically assess the epigenetic regulation of MYC we performed CUT&Tag for H3K27Ac on sample 14-00613. We observed decreased accessibility and activation at active BENC modules (Fig. 6F-G; Supplementary Fig. S8E). Additionally, these dynamic regions are bound by PU.1 and LSD1 in Kasumi-1 cells (Fig. 6F). Taken together, these data suggest regulation of BENC to decrease MYC transcription is a conserved mechanism between the cell line model and KIT-mutant AML primary cells. In sum, LSD1 and KIT inhibition is synergistically cytotoxic to patient samples with KIT mutations. With these samples, we confirm MYC target and cell cycle repression through transcriptomics and epigenetic changes.

### LSD1 and KIT inhibition results in decreased MYC expression along single cell differentiation trajectory

Recent work using single cell RNA sequencing (scRNA-seq) has revealed that AML cells show substantial heterogeneity in differentiation status, existing along a differentiation continuum(42). Cells at different relative positions on this trajectory may respond differently to treatment; however, these differences cannot be appreciated through bulk analysis. Therefore, we performed scRNA-seq to understand the transcriptional response of different cell populations within patient sample 14-00613. This analysis revealed seven transcriptionally-defined clusters (Fig. 7A; Supplementary Tables S24-S25). These clusters were classified using features of early, mid, and late maturation (Supplementary Fig. S9A-B). Given the evidence of MYC and MYC target repression as a key response to dual LSD1 and KIT inhibition, we assessed MYC expression across the myeloid trajectory (Fig. 7B-C). With combined KIT and LSD1 inhibition relative to DMSO control treatment, average MYC expression was decreased in undifferentiated cells (cluster 0) and MYC was detected in fewer cells. A similar response occurs in mid-trajectory cells (cluster 2). Taken together, dual inhibition of LSD1 and KIT in a KIT-mutant patient sample results in decreased MYC expression in immature cells. This single cell RNA-seq analysis further supports MYC regulation as a key driver of the response to dual LSD1 and KIT inhibition.

**Figure 7.**
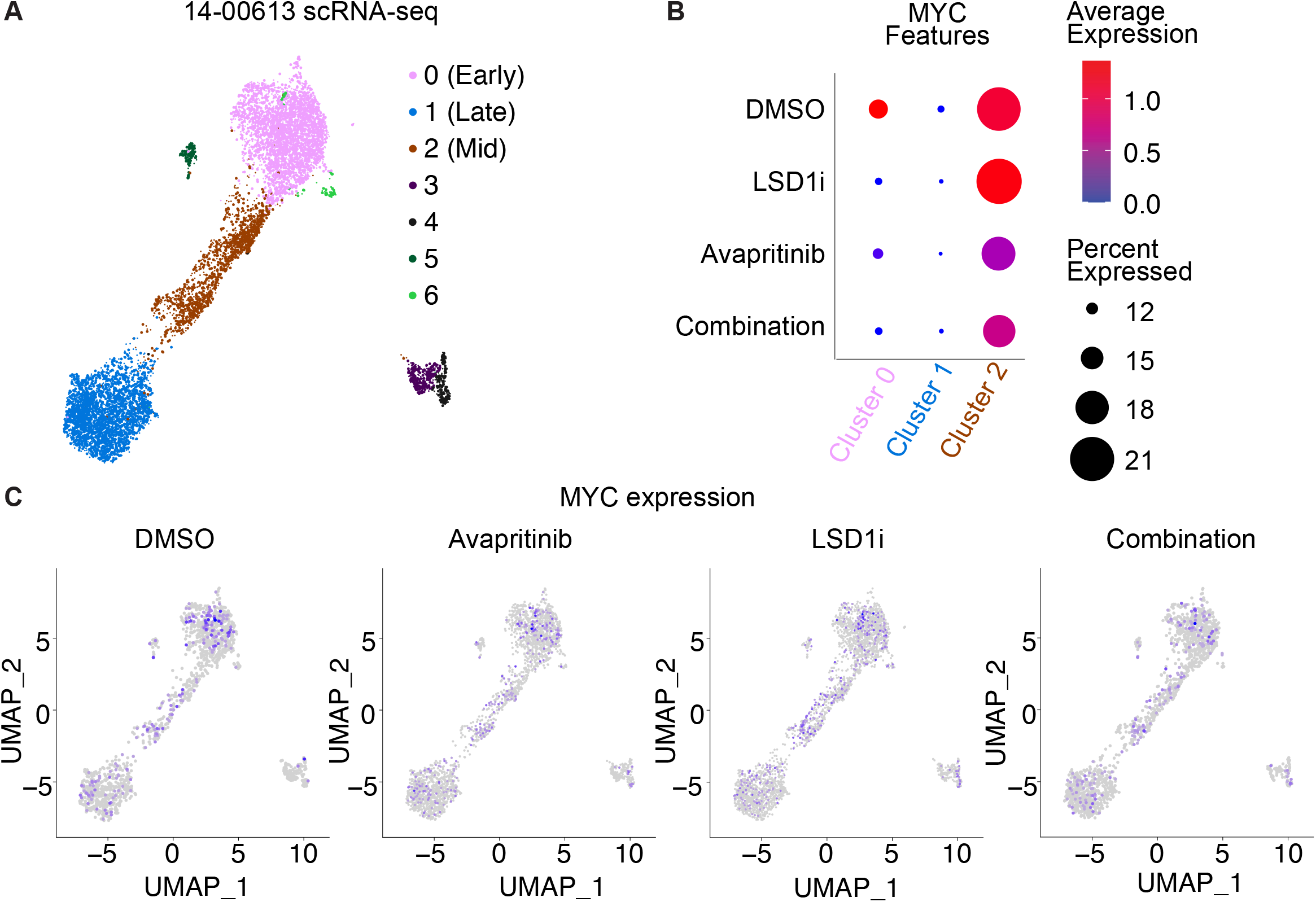
Varied degree of MYC loss along the differentiation trajectory post LSD1 and KIT inhibition. **A.** Single cell RNA-seq on patient sample 14-00613 treated with ORY-1001 (12 nM) and/or avapritinib (12 nM) for 24 h. UMAP clustering of single cell gene expression. Clustering was performed on an integrated object that included cells treated with DMSO, avapritinib, ORY-1001, or the combination. **B.** Dot plot portraying the average gene expression and percentage of cells with MYC detected per UMAP cluster in A. **C.** MYC single cell gene expression in DMSO, ORY-1001 (LSD1i), avapritinib, and the combination from single cell analysis in A.

## Discussion

This work offers mechanistic insight into the mechanism of synergy between LSD1 and KIT inhibition in KIT-mutant AML, providing rational for evaluating the combination in patient with KIT-mutant AML. The standard of care treatment for patients with KIT-mutant AML involves induction and consolidation with cytarabine-based chemotherapy(43). Following treatment, patients with CBF AML harboring mutant KIT have higher risk of relapse compared to CBF AML alone, thus the need for improved therapeutic options(44). In this study, we evaluated dual inhibition of LSD1 and KIT as a novel combination therapy for KIT-mutant AML. We show that combined inhibition of LSD1 and KIT is synergistically cytotoxic to AML cell lines harboring KIT mutations. Bulk RNA-seq revealed dysregulation of MYC and PU.1 (SPI1) transcriptional networks following combination treatment. Dual inhibition of LSD1 and KIT resulted in a loss of MYC binding and a loss of activating H3K27Ac at cell cycle genes, driving cell cycle exit. Furthermore, we identified a previously unappreciated role for PU.1 in driving MYC expression. Inhibition of LSD1 results in a global loss of PU.1 binding, including loss at the +26 kb MYC enhancer and blood enhancer cluster (BENC). This loss of PU.1 binding results in decreased enhancer acetylation and a corresponding decrease in MYC gene expression driving cell cycle exit. Dual LSD1 and KIT inhibition is also cytotoxic in AML patient samples with KIT mutations, with transcriptional and epigenetic profiling confirming the findings from cell lines. Our studies reveal that modulation of MYC and PU.1 transcriptional networks is a key feature of dual LSD1 and KIT inhibition, providing pre-clinical rational for early phase clinical trials investigating this combination in KIT-mutant AML.

PU.1 has previously been implicated as a key determinant of LSD1 inhibitor responses in MLL-rearranged AML, with LSD1 inhibition leading to activation of PU.1 bound enhancers(23,25). Consistent with this, our RNA-seq shows that PU.1 targets are activated with combined KIT and LSD1 inhibition. Interestingly, genome wide profiling of PU.1 binding revealed loss of PU.1 binding with LSD1 inhibition. Together these data suggest that PU.1 serves as a transcriptional repressor at a large number of genes. Depending on the context, PU.1 can act as a repressor or activator(45). In normal hematopoiesis PU.1 and RUNX1, a part of the core binding factor (CBF) complex, are coactivators; however, PU.1 and RUNX1-ETO, a CBF fusion, are corepressors(46). CBF fusions are expressed in the KIT-mutant AML samples used in this study. Thus, when PU.1 binding is lost due to LSD1 inhibition, it is likely reactivating genes that were repressed by PU.1/RUNX1-ETO. At other loci, our data suggest PU.1 serves as a transcriptional activator, likely through interactions with coactivators, such as histone acetyl transferases(47). In our study, PU.1 loss is accompanied by a loss of H3K27Ac at the +26 kB MYC enhancer and BENC with a corresponding decrease in MYC expression, suggesting transactivation activity. When PU.1 is knocked-down, as a model for PU.1 binding loss, we observed a decrease in acetylation at the enhancers and promoter of MYC. We hypothesize that loss of activating PU.1 complexes at these MYC enhancers leads to a loss of MYC expression and cell cycle exit.

The underlying mechanism of PU.1 binding loss genome-wide by LSD1 inhibition is not clear, but may be regulated by co-binding and post-translational modifications. We have found regions with PU.1 and LSD1 co-binding (i.e. BENC modules) and other regions where they bind independently (i.e. MYC promoter and +26 kB enhancer). In the cobound regions, PU.1 loss could be a direct result of LSD1 inhibition disrupting protein-protein interactions. However, the loss of PU.1 at regions where LSD1 is not bound, suggests an intermediate step, such as post-translational modifications. PU.1 is phosphorylated at Ser148(48,49) leading to increased activity and DNA binding. Additionally, LSD1 is known to demethylate non-histone protein substrates and could potentially regulate transcription factors, such as PU.1, in this manner. The precise mechanism of LSD1-dependent suppression of PU.1 activity in KIT-mutant AML remains an important area for future investigation.

GFI1 has been identified as a repressive transcription factor that co-binds with LSD1(23). Disruption of the LSD1-GFI1 complex has been implicated as a key mechanism of LSD1 inhibitor response in MLL-rearranged AML resulting in enhancer activation and subsequent differentiation(23). We confirmed that LSD1 inhibition also evicts GFI1 from chromatin in KIT-mutant AML, but found that this loss of GFI1 binding did not substantially alter the activity of underlying enhancers. We did identify numerous regions of differential chromatin remodeling and gene activation in response to LSD1 inhibition, arguing that additional factors are also crucial for driving drug responses. Indeed, prior studies have nominated numerous other transcription factors as key regulators of the effect of LSD1 inhibitor including CEBPA, PU.1 and MYB(23). Our results suggest that GFI1 dependent gene regulation may play a more prominent role in MLL-rearranged AML than KIT-mutant AML. It is unknown if GFI1 contributes to changes in MYC and PU.1 transcriptional networks in KIT-mutant AML, and is a key area for future study.

Collectively, our results demonstrate dual inhibition of KIT and LSD1 is synergistically cytotoxic in KIT-mutant AML cell lines and in primary patient samples. Given the propensity for KIT-mutant AML to relapse after standard of care treatment, many experts support bone marrow transplantation for such patients in first remission, although this has not yet become the standard of care. A less toxic treatment approach would be of high clinical value for patients of advanced age or with significant co-morbidities. We show here that combined KIT and LSD1 inhibition is highly effective against KIT-mutant AML, suggesting this approach could be added to standard of care treatment to decrease relapse rates or could be used in the salvage setting for individuals with relapsed disease not eligible for intensive chemotherapy. Furthermore, we identify changes in the PU.1 and MYC transcriptional networks as key biomarkers of drug efficacy. Assessment of the activity of these key transcriptional nodes could be a valuable correlate for on-target drug activity during clinical investigation. In total, our results support the investigation of combined KIT and LSD1 inhibition in early phase clinical trials for patients with KIT-mutant AML.

## Methods

### Detailed materials and methods are available in Supplementary Data

#### Cell Lines

Kasumi-1 (ATCC), CMK (DSMZ) and MOLM-13 (DSMZ) cells were cultured in RPMI (Gibco) supplemented with 20% fetal bovine serum (FBS, HyClone), 2 mM GlutaMAX (Gibco), 100 units/mL Penicillin, and 100 ug/mL Streptomycin (Gibco). K562 (ATCC) cells were cultured in RPMI supplemented with 10% FBS. SNKO-1 cells were obtained from DSMZ and cultured in RPMI with 10% FSB and 10 ng/ml GM-CSF (PeproTech). EOL-1 cells were obtained from DSMZ and cultured in RPMI with 10% FBS. All cells were cultured at 5% CO_2_ and 37°C. Cell lines were tested monthly for mycoplasma contamination.

#### Patient Samples

All patients gave informed consent to participate in this study, which had the approval and guidance of the institutional review boards at Oregon Health & Science University (OHSU), University of Utah, University of Texas Medical Center (UT Southwestern), Stanford University, University of Miami, University of Colorado, University of Florida, National Institutes of Health (NIH), Fox Chase Cancer Center and University of Kansas (KUMC). Samples were sent to the coordinating center (OHSU; IRB#9570; #4422; <u>NCT01728402</u>) where they were coded and processed. Additional processing details available in supplemental methods. Patient sample mononuclear cells were cultured in RPMI with 10% FBS and 50% HS-5 conditioned media (ATCC) or SFEMII supplemented with 1x StemSpan CD34+ Expansion Media and 1 uM UM729 (StemCell Technologies).

#### Bulk RNA-sequencing

Kasumi-1 cells were treated with 12 nM GSK-LSD1, 12 nM avapritinib, 12 nM GSK-LSD1/12 nM avapritinib or DMSO for 12 h. Total RNA was isolated using a RNeasy Plus Mini Kit (Qiagen). BGI performed the library preparation and sequencing at 50 base pair (BP) paired end (PE). For the Kasumi-1 PU.1 KD experiment, cells were treated with 1 ug/ml doxycycline, 50 nM avapritinib, 1 ug/ml doxycycline/50 nM avapritinib, or DMSO for 24 hours. Total RNA was isolated with RNeasy Plus Mini Kit (Qiagen) and libraries were prepared with Zymo-Seq RiboFree Total RNA Library Kit (Zymo). The OHSU Massively Parallel Sequencing Shared Resource (MPSSR) performed the sequencing at 100 BP single end.

#### CUT&Tag

Kasumi-1 cells were treated with 12 nM GSK-LSD1, 12 nM avapritinib, 12 nM GSK-LSD1/12 nM avapritinib or DMSO for 24 hr. Benchtop CUT&Tag was performed as previously described(50). In brief, cells were harvested and washed 2x in wash buffer. Cells were bound to magnetic beads and incubated with primary antibody overnight followed by secondary antibody for 1 h. The primary antibodies used were: H3K27Ac (ab4729, Abcam), H3K4me1 (#5326, Cell Signaling Technologies), H3K4me3 (#9751, Cell Signaling Technologies), and Normal Rabbit IgG (#2729, CST). The secondary antibody used was guinea-pig anti rabbit (Antibodies Online). Samples were incubated with pA-Tn5 and tagmented for 1 h at 37°C. DNA was extracted with phenol:chloroform extraction and amplified with custom nextera primers(51). Libraries were purified and sequenced using 37 BP PE sequencing on a NextSeq 500 sequencer (Illumina) by MPSSR.

Kasumi-1 PU.1 sh401 cells were treated with 1 ug/ml doxycycline, 50 nM avapritinib, 1 ug/ml doxycycline/50 nM avapritinib, or DMSO for 24h. 14-00613 cells were treated with 12 nM ORY-1001, 350 nM avapritinib, 12 nM ORY-1001/350 nM avapritinib, or DMSO for 24h. CUT&Tag was adapted from the benchtop version and was preformed similarly to the updated protocol(52). Nuclei were prepared and bound to magnetic beads. Samples were incubated with primary antibody overnight then secondary antibody for 30 min. pA-Tn5 (EpiCypher) was incubated with samples for 1 h, then tagmentation took place at 37°C for 1 h. DNA fragments were amplified using custom nextera primers(51). PCR products were purified and underwent 37 BP PE sequencing on a NextSeq 500 sequencer (Illumina) by MPSSR. Additional details on both CUT&Tag methods can be found in supplementary information.

#### CUT&RUN

Kasumi-1 cells were treated with 12 nM GSK-LSD1, 12 nM avapritinib, 12 nM GSK-LSD1/12 nM avapritinib or DMSO for 12h or 24 h. CUT&RUN was performed as previously described(53). In short, 500,000 cells per replicate were harvested and bound to magnetic beads. Following permeabilization, cells were incubated overnight with primary antibody. The following antibodies were used at 1:50: MYC (#13987, CST), PU.1 (MA5-15064, Invitrogen), LSD1 (ab17721, Abcam), GFI1 (ab21061, Abcam) and Normal Rabbit IgG (#2729, CST). Cells were incubated with pAG-MNase (Epicypher) in dig wash buffer then in reaction buffer. STOP buffer was used to inhibit the reaction. DNA was extracted with phenol:cholorform. Libraries were generated using NEBNext Ultra II DNA Library Prep Kit (NEB), modified for CUT&RUN as previously described(54). Libraries were sequenced on a NextSeq 500 sequencer (Illumina) using 37 BP PE sequencing by MPSSR.

#### Fluorescent Biosensor Assay

Kasumi-1 cells were transduced with ERK-TB-pHAGE-Clover-H2BmiFRP670nano and AKT-TB-pHAGE-mScarlet-H2BmiRFP670nano derived from ERK-KTR (PMID: 24949979) and AKT-KTR (PMID: 27247077) respectively. Transduced cells were sorted on a BD FACSAria Fusion housed at the OHSU Flow Cytometry Shared Resource to enrich the triple positive cells (Clover+, mScarlet+, & miRFP670nano+). For time lapse imaging, biosensor-expressed cells were resuspended in FluoroBrite DMEM (Gibco), 1x GlutaMAX (Gibco), and 20% FBS, and then seeded on a 96 well plate at density of ~10,000 cells/well. Cells were imaged on the Yokogawa CSU-X1 spinning disk confocal microscope housed in ALMC under cell culture-optimized temperature, humidity and CO2 levels. Cell images were captured every 15min for a total of 25 h after drug loading.

#### Reverse Phase Protein Array

Kasumi-1 cells were treated for 1 h and 24 h with 100nM avapritinib and 12 nM ORY-1001. Cells were washed 2x in PBS then flash frozen. Cell pellets were lysed and processed by the University of Texas MD Anderson Cancer Center Functional Proteomics RPPA Core Facility. Linear normalized values used for CausalPath and log normalized values used for heatmap generation.

#### ATAC sequencing

Patient sample 15-00807 was treated with 50 nM avapritinib, 50 nM ORY-1001, 50 nM avapritinib/50 nM ORY-1001, or DMSO for 24 hours. 50,000 cells per replicate were harvested for Fast-ATAC sequencing performed as previously described(55). Briefly, cells were tagmented with TDE1 (Illumina) for 30 min at 37°C. DNA was purified and amplified as previously described. Samples were quantified, pooled, then sequenced by Genewiz with HiSeq-X (Illumina) 75 BP PE sequencing.

#### Single Cell RNA Sequencing

Patient sample 14-00613 was treated with 12 nM ORY-1001, 12 nM avapritinib, 12 nM ORY-1001/12 nM avapritinib or DMSO for 24 hours then harvested. Cells were loaded onto the Chromium Controller (10X Genomics) according to the manufacturer’s instructions. RNA libraries were prepared per the manufacturer’s protocol, Chromium Next GEM Single Cell 3’ Reagent v3.0 Kit (Biolegend, 10X Genomics). Libraries were sequenced on a HiSeq2500 (Illumina) using 100 BP PE sequencing.

### Data Analysis

#### CUT&Tag and CUT&RUN Analysis

CUT&Tag and CUT&RUN libraries were aligned to the human genome (hg38) using Bowtie2(56) and the following options --local --very-sensitive-local --no-unal --no-mixed --no-discordant --phred33 -I 10 -X 700. Peaks were called using a custom script available upon request. High confidence peaks were defined by presence in 2/2 or 2/3 replicates. Consensus bed files were formed by merging the high confidence peaks from DMSO, avapritinib, ORY-1001, and combination using BEDTools(57). Differential peaks were identified using DESeq2(58) with default parameters. Heatmaps were produced using the ComplexHeatmap package from Bioconductor(59). Peaks were annotated to the nearest feature using ChIPseeker(60). GO analysis was performed using Genomic Regions Enrichment of Annotations Tool (GREAT)(61). Counts tables for differential peaks were produced using multicov from BEDTools(57). Counts per million (CPM) normalized tracks, global signal heatmaps, and plot profiles at specified regions were generated using DeepTools(62). The DeepTools matrix was extracted and used to assess differences in H3K27Ac signal. Active promoters were defined by the presence of H3K4me3 and H3K27Ac within 1000 bp of a TSS. Active enhancers were defined by the presence of H3K4me1 and H3K27Ac beyond 1000 bp of a TSS. CPM normalized tracks were merged with bigWigMerge(63) and visualized using Integrative Genomics Viewer(64).

#### ATAC Analysis

ATAC libraries were aligned to the human genome (hg38) using BWA-MEM(65) and sorted using SAMtools(66). Duplicates were marked with Sambamba(67), duplicates and mitochondrial reads were removed. CPM normalized tracks were generated with DeepTools(62). Differential accessibility was assessed with DESeq2(58). GO analysis was performed using GREAT(61). CPM normalized tracks were merged with bigWigMerge(63) and visualized using Integrative Genomics Viewer(64).

#### Bulk RNA Sequencing Analysis

Raw reads were trimmed with Trimmomatic(68) and aligned with STAR(69). Differential expression analysis was performed using DESeq2(58). Raw p values were adjusted for multiple comparisons using the Benjamini-Hochberg method. GO analysis was performed using Enrichr(70) and Gene Set Enrichment Analysis (GSEA)(71).

#### Single Cell RNA Sequencing Analysis

Sequencing output from 10X mRNA sequencing libraries were aligned to human genome (hg38) using CellRanger. Filtered feature matrices were analyzed using Seurat(72). Cells expressing greater than 20% mitochondrial RNA were excluded as non-viable. Data integration revealed seven transcriptional clusters. Cluster identity was established via expression of early (CD34, SOX4, ERG, MYB, and GATA2), mid (CEBPD), and late (LYZ, ITGAX, CD14) features of hematopoiesis.

#### Fluorescent Biosensor Analysis

Cells were tracked and subcellular fluorescence signals were measured using IMARIS software at ALMC. Heatmaps and lineplots of AKT and ERK activity were created using customized Python algorithm and Excel. Cells were separated into different AKT dynamics clusters by hierarchical clustering. Cell number was scaled to be equal across conditions and the abundance of differently treated cells per each cluster was calculated.

#### Quantification and Statistical Analysis

Data are shown as mean ± SEM. Prism software (version 9.1; Prism Software Corp.) or RStudio was used to perform statistical analysis, which are described in the figure legends. All data were analyzed with Kolmogorov-Smirnov (KS) test or ANOVA followed by Holm-Sidak correction. For differential analysis of RNA-seq, CUT&Tag, CUT&RUN, and ATAC-seq P values were adjusted for repeated testing with a false discovery rate using the Benjamini-Hochberg method(73).

### Data Availability

Data generated in this study are available within the article and its supplementary data files.

## Acknowledgements

We thank the following OHSU core facilities for their assistance: Advanced Light Microscopy, Flow Cytometry Shared Resource, Massive Parallel Sequencing Shared Resource, ExaCloud Cluster Computational Resource, and the Advanced Computing Center. Funding was provided by an American Society of Hematology Research Restart Award, an American Society of Hematology Scholar Award and 1 K08 CA245224 from NCI awarded to T.P.B. The Functional Proteomics RPPA Core is supported by MD Anderson Cancer Center Support Grant # 5 P30 CA016672-40.

## Author contributions

B.M.S., T.P.B., D.J.C., B.J.D., and J.E.M. designed research; B.M.S., Y.H.T., and T.P.B. performed research; B.M.S., T.P.B., J.V., G.L.K., W.Y., W.H., D.J.C., J.E., Y.H.T. and T.A.L. contributed new reagents/analytic tools; B.M.S., Y.H.T., G.B.M., T.P.B., B.J.D., and J.E.M. analyzed data; B.M.S wrote manuscript; B.M.S, T.P.B., and J.E.M. reviewed and edited the manuscript.

## References

1. Kelly LM, Gilliland DG. Genetics of myeloid leukemias. Annu Rev Genomics Hum Genet. 2002 Sep 1;3(1):179–98.

2. Scheijen B, Griffin JD. Tyrosine kinase oncogenes in normal hematopoiesis and hematological disease. Oncogene. 2002 May;21(21):3314–33.

3. Gilliland DG, Griffin JD. The roles of FLT3 in hematopoiesis and leukemia. Blood. 2002 Sep 1;100(5):1532–42.

4. Bullinger L, Döhner K, Döhner H. Genomics of Acute Myeloid Leukemia Diagnosis and Pathways. J Clin Oncol. 2017 Feb 13;35(9):934–46.

5. Corces-Zimmerman MR, Hong W-J, Weissman IL, Medeiros BC, Majeti R. Preleukemic mutations in human acute myeloid leukemia affect epigenetic regulators and persist in remission. Proc Natl Acad Sci. 2014 Feb 18;111(7):2548–53.

6. Ueyama J, Kure A, Okuno K, Sano H, Tamoto N, Kanzaki S. [Treatment with a tyrosine-kinase inhibitor of for c-KIT mutation and AML1-ETO double positive refractory acute myeloid leukemia]. Rinshō Ketsueki Jpn J Clin Hematol. 2012;53(4):460–4.

7. Cairoli R, Beghini A, Morello E, Grillo G, Montillo M, Larizza L, et al. Imatinib mesylate in the treatment of Core Binding Factor leukemias with KIT mutations: A report of three cases. Leuk Res. 2005 Apr 1;29(4):397–400.

8. Megías-Vericat JE, Ballesta-López O, Barragán E, Martínez-Cuadrón D, Montesinos P. Tyrosine kinase inhibitors for acute myeloid leukemia: A step toward disease control? Blood Rev. 2020 Nov 1;44:100675.

9. Bewersdorf JP, Shallis R, Stahl M, Zeidan AM. Epigenetic therapy combinations in acute myeloid leukemia: what are the options? Ther Adv Hematol. 2019 Jan 1;10:2040620718816698.

10. Salamero O, Montesinos P, Willekens C, Pérez-Simón JA, Pigneux A, Récher C, et al. First-in-Human Phase I Study of Iadademstat (ORY-1001): A First-in-Class Lysine-Specific Histone Demethylase 1A Inhibitor, in Relapsed or Refractory Acute Myeloid Leukemia. J Clin Oncol. 2020 Dec 20;38(36):4260–73.

11. Odenike O, Wolff JE, Borthakur G, Aldoss IT, Rizzieri D, Prebet T, et al. Results from the first-in-human study of mivebresib (ABBV-075), a pan-inhibitor of bromodomain and extra terminal proteins, in patients with relapsed/refractory acute myeloid leukemia. J Clin Oncol. 2019 May 20;37(15_suppl):7030–7030.

12. Dombret H, Preudhomme C, Berthon C, Raffoux E, Thomas X, Vey N, et al. A Phase 1 Study of the BET-Bromodomain Inhibitor OTX015 in Patients with Advanced Acute Leukemia. Blood. 2014 Dec 6;124(21):117–117.

13. Braun TP, Coblentz C, Smith BM, Coleman DJ, Schonrock Z, Carratt SA, et al. Combined inhibition of JAK/STAT pathway and lysine-specific demethylase 1 as a therapeutic strategy in CSF3R/CEBPA mutant acute myeloid leukemia. Proc Natl Acad Sci. 2020 Jun 16;117(24):13670–9.

14. Avellino R, Delwel R. Expression and regulation of C/EBPα in normal myelopoiesis and in malignant transformation. Blood. 2017 Apr 13;129(15):2083–91.

15. Speck NA, Gilliland DG. Core-binding factors in haematopoiesis and leukaemia. Nat Rev Cancer. 2002 Jul;2(7):502–13.

16. Paschka P, Marcucci G, Ruppert AS, Mrózek K, Chen H, Kittles RA, et al. Adverse Prognostic Significance of KIT Mutations in Adult Acute Myeloid Leukemia With inv(16) and t(8;21): A Cancer and Leukemia Group B Study. J Clin Oncol. 2006 Aug 20;24(24):3904–11.

17. Brizzi MF, Dentelli P, Rosso A, Yarden Y, Pegoraro L. STAT Protein Recruitment and Activation in c-Kit Deletion Mutants. J Biol Chem. 1999 Jun 11;274(24):16965–72.

18. Larizza L, Magnani I, Beghini A. The Kasumi-1 cell line: a t(8;21)-kit mutant model for acute myeloid leukemia. Leuk Lymphoma. 2005 Jan;46(2):247–55.

19. Lück SC, Russ AC, Du J, Gaidzik V, Schlenk RF, Pollack JR, et al. KIT mutations confer a distinct gene expression signature in core binding factor leukaemia. Br J Haematol. 2010;148(6):925–37.

20. Evans EK, Gardino AK, Kim JL, Hodous BL, Shutes A, Davis A, et al. A precision therapy against cancers driven by KIT/PDGFRA mutations. Sci Transl Med. 2017 Nov 1;9(414):eaao1690.

21. Kerenyi MA, Shao Z, Hsu Y-J, Guo G, Luc S, O’Brien K, et al. Histone demethylase Lsd1 represses hematopoietic stem and progenitor cell signatures during blood cell maturation. Morrison SJ, editor. eLife. 2013 Jun 18;2:e00633.

22. Harris WJ, Huang X, Lynch JT, Spencer GJ, Hitchin JR, Li Y, et al. The histone demethylase KDM1A sustains the oncogenic potential of MLL-AF9 leukemia stem cells. Cancer Cell. 2012 Apr 17;21(4):473–87.

23. Maiques-Diaz A, Spencer GJ, Lynch JT, Ciceri F, Williams EL, Amaral FMR, et al. Enhancer Activation by Pharmacologic Displacement of LSD1 from GFI1 Induces Differentiation in Acute Myeloid Leukemia. Cell Rep. 2018 Mar 27;22(13):3641–59.

24. McGrath JP, Williamson KE, Balasubramanian S, Odate S, Arora S, Hatton C, et al. Pharmacological Inhibition of the Histone Lysine Demethylase KDM1A Suppresses the Growth of Multiple Acute Myeloid Leukemia Subtypes. Cancer Res. 2016 Apr 1;76(7):1975–88.

25. Cusan M, Cai SF, Mohammad HP, Krivtsov A, Chramiec A, Loizou E, et al. LSD1 inhibition exerts its antileukemic effect by recommissioning PU.1- and C/EBPα-dependent enhancers in AML. Blood. 2018 12;131(15):1730–42.

26. Barth J, Abou-El-Ardat K, Dalic D, Kurrle N, Maier A-M, Mohr S, et al. LSD1 inhibition by tranylcypromine derivatives interferes with GFI1-mediated repression of PU.1 target genes and induces differentiation in AML. Leukemia. 2019 Jun;33(6):1411–26.

27. Moreau-Gachelin F. Spi-1/PU.1: an oncogene of the Ets family. Biochim Biophys Acta BBA - Rev Cancer. 1994 Dec 30;1198(2):149–63.

28. Eilers M, Eisenman RN. Myc’s broad reach. Genes Dev. 2008 Oct 15;22(20):2755–66.

29. Bahr C, von Paleske L, Uslu VV, Remeseiro S, Takayama N, Ng SW, et al. A Myc enhancer cluster regulates normal and leukaemic haematopoietic stem cell hierarchies. Nature. 2018 Jan;553(7689):515–20.

30. Shi J, Whyte WA, Zepeda-Mendoza CJ, Milazzo JP, Shen C, Roe J-S, et al. Role of SWI/SNF in acute leukemia maintenance and enhancer-mediated Myc regulation. Genes Dev. 2013 Dec 15;27(24):2648–62.

31. Fishilevich S, Nudel R, Rappaport N, Hadar R, Plaschkes I, Iny Stein T, et al. GeneHancer: genome-wide integration of enhancers and target genes in GeneCards. Database J Biol Databases Curation [Internet]. 2017 Apr 17 [cited 2021 Jun 16];2017. Available from: https://www.ncbi.nlm.nih.gov/pmc/articles/PMC5467550/

32. Sears R. The Life Cycle of C-Myc: From Synthesis to Degradation. Cell Cycle Georget Tex. 2004 Oct 1;3:1133–7.

33. Gregory MA, Qi Y, Hann SR. Phosphorylation by Glycogen Synthase Kinase-3 Controls c-Myc Proteolysis and Subnuclear Localization*. J Biol Chem. 2003 Dec 19;278(51):51606–12.

34. Manning BD, Toker A. AKT/PKB Signaling: Navigating the Network. Cell. 2017 Apr;169(3):381–405.

35. Darici S, Alkhaldi H, Horne G, Jørgensen HG, Marmiroli S, Huang X. Targeting PI3K/Akt/mTOR in AML: Rationale and Clinical Evidence. J Clin Med. 2020 Sep 11;9(9).

36. Peck B, Ferber EC, Schulze A. Antagonism between FOXO and MYC Regulates Cellular Powerhouse. Front Oncol. 2013 Apr 25;3:96.

37. Sears R, Nuckolls F, Haura E, Taya Y, Tamai K, Nevins JR. Multiple Ras-dependent phosphorylation pathways regulate Myc protein stability. Genes Dev. 2000 Oct 1;14(19):2501–14.

38. Zhou A, Lin K, Zhang S, Chen Y, Zhang N, Xue J, et al. Nuclear GSK3β promotes tumorigenesis by phosphorylating KDM1A and inducing its deubiquitylation by USP22. Nat Cell Biol. 2016 Sep;18(9):954–66.

39. Tyner JW, Tognon CE, Bottomly D, Wilmot B, Kurtz SE, Savage SL, et al. Functional genomic landscape of acute myeloid leukaemia. Nature. 2018 Oct;562(7728):526–31.

40. Buenrostro JD, Wu B, Chang HY, Greenleaf WJ. ATAC-seq: A Method for Assaying Chromatin Accessibility Genome-Wide. Curr Protoc Mol Biol. 2015;109(1):21.29.1–21.29.9.

41. cdc2–cyclin B regulates eEF2 kinase activity in a cell cycle- and amino acid-dependent manner. EMBO J. 2008 Apr 9;27(7):1005–16.

42. van Galen P, Hovestadt V, Wadsworth II MH, Hughes TK, Griffin GK, Battaglia S, et al. Single-Cell RNA-Seq Reveals AML Hierarchies Relevant to Disease Progression and Immunity. Cell. 2019 Mar 7;176(6):1265–1281.e24.

43. Zhu H-H, Zhang X-H, Qin Y-Z, Liu D-H, Jiang H, Chen H, et al. MRD-directed risk stratification treatment may improve outcomes of t(8;21) AML in the first complete remission: results from the AML05 multicenter trial. Blood. 2013 May 16;121(20):4056–62.

44. Tarlock K, Alonzo TA, Wang Y-C, Gerbing RB, Ries R, Loken MR, et al. Functional Properties of KIT Mutations Are Associated with Differential Clinical Outcomes and Response to Targeted Therapeutics in CBF Acute Myeloid Leukemia. Clin Cancer Res. 2019 Aug 15;25(16):5038–48.

45. van Riel B, Rosenbauer F. Epigenetic control of hematopoiesis: the PU.1 chromatin connection. Biol Chem. 2014 Nov 1;395(11):1265–74.

46. Hu Z, Gu X, Baraoidan K, Ibanez V, Sharma A, Kadkol S, et al. RUNX1 regulates corepressor interactions of PU.1. Blood. 2011 Jun 16;117(24):6498–508.

47. Bai Y, Srinivasan L, Perkins L, Atchison ML. Protein Acetylation Regulates Both PU.1 Transactivation and Igκ 3′ Enhancer Activity. J Immunol. 2005 Oct 15;175(8):5160–9.

48. Pongubala JMR, Van Beveren C, Nagulapalli S, Klemsz MJ, McKercher SR, Maki RA, et al. Effect of PU.1 Phosphorylation on Interaction with NF-EM5 and Transcriptional Activation. Science. 1993;259(5101):1622–5.

49. Rieske P, Pongubala JR. AKT Induces Transcriptional Activity of PU.1 through Phosphorylation-mediated Modifications within Its Transactivation Domain *. J Biol Chem. 2001 Mar 1;276(11):8460–8.

50. Kaya-Okur HS, Wu SJ, Codomo CA, Pledger ES, Bryson TD, Henikoff JG, et al. CUT&Tag for efficient epigenomic profiling of small samples and single cells. Nat Commun. 2019 Apr 29;10(1):1–10.

51. Buenrostro JD, Wu B, Litzenburger UM, Ruff D, Gonzales ML, Snyder MP, et al. Single-cell chromatin accessibility reveals principles of regulatory variation. Nature. 2015 Jul;523(7561):486–90.

52. Henikoff S, Henikoff JG, Kaya-Okur HS, Ahmad K. Efficient chromatin accessibility mapping in situ by nucleosome-tethered tagmentation. Bonasio R, Tyler JK, Danko CG, Cao J, editors. eLife. 2020 Nov 16;9:e63274.

53. Skene PJ, Henikoff S. An efficient targeted nuclease strategy for high-resolution mapping of DNA binding sites. Reinberg D, editor. eLife. 2017 Jan 12;6:e21856.

54. Liu N, Hargreaves VV, Zhu Q, Kurland JV, Hong J, Kim W, et al. Direct Promoter Repression by BCL11A Controls the Fetal to Adult Hemoglobin Switch. Cell. 2018 Apr 5;173(2):430–442.e17.

55. Corces MR, Buenrostro JD, Wu B, Greenside PG, Chan SM, Koenig JL, et al. Lineage-specific and single-cell chromatin accessibility charts human hematopoiesis and leukemia evolution. Nat Genet. 2016 Oct;48(10):1193–203.

56. Langmead B, Salzberg SL. Fast gapped-read alignment with Bowtie 2. Nat Methods. 2012 Apr;9(4):357–9.

57. Quinlan AR, Hall IM. BEDTools: a flexible suite of utilities for comparing genomic features. Bioinformatics. 2010 Mar 15;26(6):841–2.

58. Love MI, Huber W, Anders S. Moderated estimation of fold change and dispersion for RNA-seq data with DESeq2. Genome Biol. 2014 Dec 5;15(12):550.

59. Gu Z, Eils R, Schlesner M. Complex heatmaps reveal patterns and correlations in multidimensional genomic data. Bioinformatics. 2016 Sep 15;32(18):2847–9.

60. Yu G, Wang L-G, He Q-Y. ChIPseeker: an R/Bioconductor package for ChIP peak annotation, comparison and visualization. Bioinformatics. 2015 Jul 15;31(14):2382–3.

61. McLean CY, Bristor D, Hiller M, Clarke SL, Schaar BT, Lowe CB, et al. GREAT improves functional interpretation of cis-regulatory regions. Nat Biotechnol. 2010 May;28(5):495–501.

62. Ramírez F, Ryan DP, Grüning B, Bhardwaj V, Kilpert F, Richter AS, et al. deepTools2: a next generation web server for deep-sequencing data analysis. Nucleic Acids Res. 2016 Jul 8;44(W1):W160–5.

63. Kuhn RM, Haussler D, Kent WJ. The UCSC genome browser and associated tools. Brief Bioinform. 2013 Mar 1;14(2):144–61.

64. Robinson JT, Thorvaldsdóttir H, Winckler W, Guttman M, Lander ES, Getz G, et al. Integrative Genomics Viewer. Nat Biotechnol. 2011 Jan;29(1):24–6.

65. Li H, Durbin R. Fast and accurate short read alignment with Burrows–Wheeler transform. Bioinformatics. 2009 Jul 15;25(14):1754–60.

66. Li H, Handsaker B, Wysoker A, Fennell T, Ruan J, Homer N, et al. The Sequence Alignment/Map format and SAMtools. Bioinformatics. 2009 Aug 15;25(16):2078–9.

67. Tarasov A, Vilella AJ, Cuppen E, Nijman IJ, Prins P. Sambamba: fast processing of NGS alignment formats. Bioinformatics. 2015 Jun 15;31(12):2032–4.

68. Bolger AM, Lohse M, Usadel B. Trimmomatic: a flexible trimmer for Illumina sequence data. Bioinformatics. 2014 Aug 1;30(15):2114–20.

69. Dobin A, Davis CA, Schlesinger F, Drenkow J, Zaleski C, Jha S, et al. STAR: ultra-fast universal RNA-seq aligner. Bioinformatics. 2013 Jan 1;29(1):15–21.

70. Kuleshov MV, Jones MR, Rouillard AD, Fernandez NF, Duan Q, Wang Z, et al. Enrichr: a comprehensive gene set enrichment analysis web server 2016 update. Nucleic Acids Res. 2016 Jul 8;44(W1):W90–7.

71. Subramanian A, Tamayo P, Mootha VK, Mukherjee S, Ebert BL, Gillette MA, et al. Gene set enrichment analysis: A knowledge-based approach for interpreting genome-wide expression profiles. Proc Natl Acad Sci. 2005 Oct 25;102(43):15545–50.

72. Butler A, Hoffman P, Smibert P, Papalexi E, Satija R. Integrating single-cell transcriptomic data across different conditions, technologies, and species. Nat Biotechnol. 2018 May;36(5):411–20.

73. Johnson WE, Li C, Rabinovic A. Adjusting batch effects in microarray expression data using empirical Bayes methods. Biostatistics. 2007 Jan 1;8(1):118–27.

74. Ianevski A, Giri AK, Aittokallio T. SynergyFinder 2.0: visual analytics of multi-drug combination synergies. Nucleic Acids Res. 2020 Jul 2;48(W1):W488–93.

