## Supplementary Materials for "PU.1 and MYC transcriptional network defines synergistic drug responses to KIT and LSD1 inhibition in acute myeloid leukemia"

### Supplementary Methods

*Western Blots*

Kasumi-1 cells were treated with 12 nM ORY-1001, 12 nM avapritinib, 12 nM ORY-1001/12 nM avapritinib or DMSO for 24 h. Cells were lysed in 1x SDS sample buffer (25 mmol/L Tris (pH 6.8), 1% SDS, 5% glycerol, 3% β-mercaptoethanol, and 0.03% bromophenol blue) with sonication (50% amplitude, 2 – 15 second pulses). Samples were spun at 14,000 x g for 10 minutes at 4°C then incubated for 5 minutes at 95°C. Samples were run on Criterion 4–15% Tris-HCl gradient gels (Bio-Rad). Gels were transferred to PVDF membranes, then blocked in Tris-buffered saline with 0.05% Tween (TBST) and 5% milk. Blots were probed with MYC (#13987, CST), PU.1 (MA5-15064, Invitrogen), or β-actin (#8457, CST) antibodies at 1:1000 in TBST overnight at 4°C. Membranes were incubated for 1 hour at room temperature with HRP-conjugated secondary antibody (CST, 7074S) at 1:5000. SuperSignalTM West Pico PLUS Chemiluminescent Substrate (ThermoFisher) was used to develop blots. Imaging was done on BioRad ChemiDoc Imaging System and analyzed using Image Lab (BioRad).

*CUT&Tag*

Benchtop CUT&Tag was performed as previously described (50). Cells were counted, harvested, and centrifuged for 5 min at 300xg at room temperature. Cells were washed 2X in 1.5 mL wash buffer (20 mM HEPES pH 7.5, 150 mM NaCl, 0.5 mM Spermidine, 1× Protease inhibitor cocktail). Concanavalin A magnetic coated beads (Bangs Laboratories) were activated in binding buffer by washing 2X (20 mM HEPES pH 7.5, 10 mM KCl, 1 mM CaCl2, 1 mM MnCl2). Washed cells were separated into 100,000 cell aliquots and 10 ul of activated beads were added to each sample. Samples rotated end over end for 7 minutes at room temperature. A magnetic stand was used to separate beads, supernatant was removed. Primary antibody was diluted 1:50 in antibody buffer (20 mM HEPES pH 7.5, 150mM NaCl, 0.5 mM Spermidine, 1× Protease inhibitor cocktail, 0.05% digitonin, 2 mM EDTA, 0.1% BSA). Cells were incubated overnight at 4°C on a nutator. Primary antibody was replaced with a guinea-pig anti rabbit secondary antibody diluted to 1:100 in wash buffer (Antibodies Online). Samples were incubated for 45 minutes at room temperature on nutator. Secondary antibody was removed and samples were washed 2X in dig-wash buffer (20 mM HEPES pH 7.5, 150 mM NaCl, 0.5 mM Spermidine, 1× Protease inhibitor cocktail, 0.05% Digitonin). pA-Tn5 transposase, prepared and loaded with adaptors as previously described(21), was diluted 1:100 in dig-300 buffer (20 mM HEPES pH 7.5, 300 mM NaCl, 0.5 mM Spermidine, 1× Protease inhibitor cocktail, 0.01% digitonin) and added to samples. Samples were incubated for 1 hour at room temperature on nutator. Samples were washed 2X with dig-300 buffer then resuspended in tagmentation buffer (dig-300 buffer with 1 mM MgCl2). Samples were incubated at 37°C for 1 hour. DNA was extracted with phenol:chloroform extraction. Samples were amplified by PCR using custom nextera primers at 400 nM and NEBNext HiFi 2x PCR Master Mix (New England Biolabs)(51). PCR conditions were set to: 72°C for 5 minutes, 98°C for 30 seconds, 14 cycles of 98°C for 10 sec, 63°C for 10 sec, and 72°C for 1 minute. Libraries were purified with AMPure Beads (Beckman) and sequenced on a NextSeq 500 sequencer (Illumina) using 37 BP PE sequencing by MPSSR.

Benchtop CUT&Tag protocol was updated and performed as described(52). Nuclei were prepared by incubating cell on ice for 10 minutes with NE1 buffer (20 mM HEPES-KOH pH 7.9, 10 mM KCl, 0.5 mM spermidine, 0.1% Triton X-100, 10% glycerol, 1x Protease inhibitor cocktail). Cells were centrifuged at 1300xg for 4 minutes at 4°C and supernatant was removed. Nuclei extract was resuspended in wash buffer (20 mM HEPES pH 7.5, 150 mM NaCl, 0.5 mM Spermidine, 1× Protease inhibitor cocktail) and 10% DMSO and frozen at -80°C. Briefly, 100,000 nuclei were thawed and incubated with 5 ul activated Concanavalin A beads (Bangs Laboratory) for 10 minutes at room temperature. Supernatant was removed and beads were resuspended in wash buffer. Samples were incubated with primary antibody diluted 1:100 in antibody buffer (wash buffer with 2 mM EDTA, 0.1% BSA) overnight at 4°C on the nutator. Primary antibody was removed, and beads incubated with guinea-pig anti rabbit secondary antibody diluted to 1:100 in wash buffer (Antibodies Online) for 30 minutes on nutator at room temperature. Beads were washed 2x with wash buffer. Beads incubated

with pA-Tn5 (EpiCypher) diluted 1:20 in 300-wash buffer (20 mM HEPES pH 7.5, 300 mM NaCl, 0.5 mM Spermidine, 1× Protease inhibitor cocktail) for 1 h at room temperature on the nutator. Beads were washed 2x in 300-wash buffer and resuspended in tagmentation solution (300-wash buffer with 10 mM $MgCl_2$). Samples incubated at 37°C for 1 hour in a PCR cycler with heated lid then were washed 1x in TAPS wash buffer (10 mM TAPS, 0.2 mM EDTA). Beads were resuspended in 0.1% SDS release solution (0.1% SDS, 10 mM TAPS) and incubated for 1 hour at 58°C in a PCR cycler with heated lid. Triton neutralization solution (0.67% triton-X100) was added to bead slurry. Samples were amplified by PCR using custom nextera primers at 400 nM and NEBNext HiFi 2x PCR Master Mix (New England Biolabs)(24). PCR conditions were set to: 58°C for 5 minutes, 72°C for 5 minutes, 98°C for 30 seconds, 12 cycles of 98°C for 10 sec, 60°C for 10 sec, and 72°C for 1 minute. Libraries were purified with HighPrep PCR Clean-up System (MagBio) and sequenced on a NextSeq 500 sequencer (Illumina) using 37 BP PE sequencing by MPSSR.

*CUT&RUN*

CUT&RUN was performed as described previously(53). Briefly, concanavalin A magnetic coated beads (Bangs Laboratories) were washed 2x in binding buffer (20 mM HEPES pH 7.5, 10 mM KCl, 1 mM $CaCl_2$, 1 mM $MnCl_2$) to activate. 500,000 cells per replicate were washed 2x with wash buffer (20 mM HEPES pH 7.5, 150 mM NaCl, 0.5 mM Spermidine, 1× Protease inhibitor cocktail). Cells were bound to beads by nutating for 10 minutes at room temperature. Cells were permeabilized and incubated over night at 4°C on nutator with primary antibody in antibody buffer (wash buffer, 0.001% digitonin, 3 mM EDTA). Bead slurry was washed 2x with dig wash buffer (wash buffer, 0.001% dig) and resuspended with dig wash buffer and 1x pAG-MNase (Epicypher). Cell were incubated for 10 minutes on nutator at room temperature then washed 2x with dig wash buffer followed by resuspension in pAG-MNase reaction mix (dig wash buffer, 2 mM $CaCl_2$). Bead slurry was incubated for 2 hours at 4°C on nutator. STOP buffer (340 mM NaCl, 20 mM EDTA, 4 mM EGTA, 50 ug/mL RNase A, 50 ug/mL glycogen, 0.02% dig) was then added, then tubes were incubated at 37°C for 10 minutes. DNA was extracted using phenol:cholorform extraction. Libraries were prepared using NEBNext Ultra II DNA Library Prep Kit (NEB), modified for CUT&RUN as previously described (54). After adapter ligation fragments were cleaned up with 1.75x AMPure beads (Beckman). Following PCR amplification, libraries were purified 2x with 1.2x AMPure beads to rid of adaptor fragments. Libraries were quantified on the 2100 Bioanalyzer instrument (Agilent) with the High Sensitivity DNA Analysis Kit (Agilent). Libraries were pooled and sequenced by MPSSR on a NextSeq 500 sequencer (Illumina) using 37 BP PE sequencing.

*ATAC sequencing*

Samples were prepared as previously described(55). In brief, cells were resuspended in cold PBS and tagmentation master mix (25 ul of 2× tagmentation buffer, 2.5 ul of TDE1 [Illumina], 0.5 ul of 1% digitonin; 2x tagmentation buffer: 66 mM Tris-Acetate, pH 7.8, 132 mM potassium acetate, 20 mM magnesium acetate, 32% v/v N,N-Dimethylformamide) was added. Samples were incubated at 37°C for 30 minutes. DNA was purified using Zymo Clean and Concentrator 5 Kit (Zymo). Transposed DNA was amplified and purified as described previously with adapted primers(40,51). Samples were quantified using Qubit dsDNA HS Assay Kit (Invitrogen), pooled, and sequenced by Genewiz with a HiSeq-X (Illumina) using 75 BP PE sequencing.

*Patient Samples*

Mononuclear cells were isolated by Ficoll gradient centrifugation from freshly obtained bone marrow aspirates or peripheral blood draws. Clinical, prognostic, genetic, cytogenetic, and pathologic lab values as well as treatment and outcome data were manually curated from patient electronic medical records. Genetic characterization of the leukemia samples included results of a clinical deep-sequencing panel of genes commonly mutated in hematologic malignancies (Sequenome and GeneTrails (OHSU); Foundation Medicine (UTSW); Genoptix; and Illumina).

*Histology*

Prior to staining, Kasumi-1 cells were treated with avapritinib (12 nM; Selleck) and/or ORY-1001 (12 nM; Selleck) for 72 h. Cells were prepared for CytoSpin 3 Shandon as described by manufacturer's protocol (ThermoShandon). Slides were fixed and stained using May-Grunwald Giemsa. Samples were imaged using Zeiss ApoTome housed in the Advanced Light Microscopy Core (ALMC) at OHSU.

*Colony assay*

Whole bone marrow was obtained (AllCells) and CD34+ cells were selected using CD34 MicroBead Kit (Miltenyi Biotec) according to manufacturer's instructions. For the colony assay, 2000 CD34+ cells were used per replicate and plated in MethoCultTM H4435 Enriched (StemCell Technologies). The four groups were treated with avapritinib (12 nM), ORY-1001 (12 nM), avapritinib and ORY-1001 (12 nM/drug), or DMSO. Plates were incubated for 14 days in 5% CO2 and 37°C. Samples were imaged using STEMvision (StemCell Technologies) and blinded prior to counting by another investigator by assigning letters randomly. ImageJ (NIH) was used to count colonies after blinding.

*Drug Synergy*

Drug synergy was assessed using an 8 x 8 matrix of drug concentrations. Cells were treated for 72 hours prior to MTS assay to evaluate viability. Cell viability was used to calculate drug synergy with SynergyFinder based on the ZIP reference model(74).

*Flow Cytometry*

To assess myeloid differentiation, Kasumi-1 cells were treated with 12 nM ORY-1001, 12 nM avapritinib, 12 nM ORY-1001/12 nM avapritinib or DMSO for 72 h. PerCP-e710 CD86 (Invitrogen) was used per the manufacture's protocol. To assess apoptosis, Kasumi-1 cells were treated with 12 nM ORY-1001, 36 nM avapritinib, 12 nM ORY-1001/36 nM avapritinib or DMSO for 48 h. APC Annexin V apoptosis detection kit (Invitrogen) was used according to the manufacture's protocol. For cell cycle analysis, Kasumi-1 cells were treated with 12 nM ORY-1001, 36 nM avapritinib, 12 nM ORY-1001/12 nM avapritinib or DMSO for 24, 48, and 72 h. Cells were fixed using ethyl alcohol and stained with propidium iodide (BioLegend). Strained cells were analyzed on a LSRFortessa flow cytometer (BD) housed in the OHSU Flow Cytometry Shared Resource.

*RNA Interference*

Two SMARTvector Inducible Human SPI1 hEF1a-TurboRFP shRNAs for SPI1/PU.1 were obtained from Horizon Discovery (230493625, 230705089). Lentivirus was produced by transfecting Lenti-X 293T cells (Clontech) with the SMARTvector transfer plasmid and packaging/pseudotyping plasmids. psPAX2 was a gift from Didier Trono (Addgene plasmid #12260; http://n2t.net/addgene:12260 ; RRID:Addgene_12260) and pMD2.G was a gift from Didier Trono (Addgene plasmid #12259 ; http://n2t.net/addgene:12259 ; RRID:Addgene_12259). The supernatants containing lentivirus was collected after 48 hours of culture and filtered with a 0.45 um filter. Kasumi-1 cells were transduced with virus via spinnoculation in the presence of polybrene. Transduced cells were selected with 1 ug/ml puromycin to produce a stable cell line.

### Supplementary Tables

Supplementary Tables S1-S8: Supplementary tables for cell lines

Supplementary Tables S9-S16: Supplementary tables for cell lines

Supplementary Tables S17-S25: Supplementary tables for primary patient samples

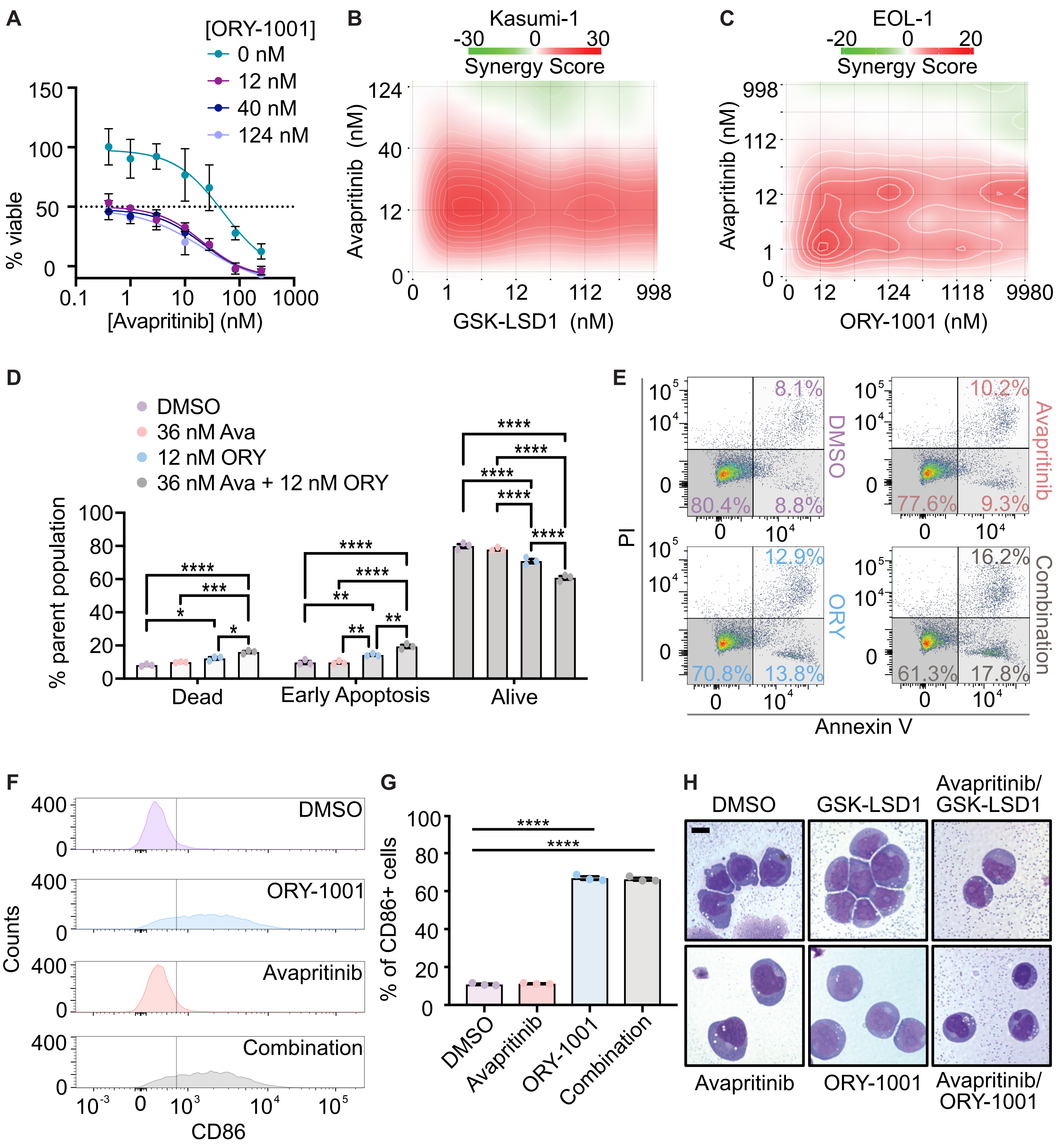


**Supplementary Figure 1. Characterization of specificity and response of KIT and LSD1 inhibition**

**A.** Viability assessment for Kasumi-1 cells treated for 72 h with avapritinib and ORY-1001 at different concentrations. Values normalized to average viability of cells with highest dose of both drugs. **B, C.** Drug matrix of Kasumi-1 or EOL-1 cells treated for 72 h with avapritinib and GSK-LSD1 or ORY-1001, respectively, with synergy assessed by ZIP score. **D, E**. Assessment of apoptosis flow cytometry of Kasumi-1 cells treated for 48 h with avapritinib (36 nM) and/or ORY-1001 (12 nM) as a percent of the parent population and representative plots of PI and Annexin V for each condition; two-way ANOVA with Holm-Sidak correction. **F, G.** Flow cytometric assessment of CD86 of Kasumi-1 cells treated for 72 h with avapritinib (12 nM) and/or ORY-1001 (12 nM). Representative histogram of CD86 signal with gray line representing CD86 positive gate and bar graph of CD86 signal for all replicates; one-way ANOVA with Holm-Sidak correction. **H.** Cytospin of Kasumi-1 cells treated for 72 h with avapritinib (12 nM), ORY-1001 (12 nM), GSK-LSD1 (12 nM), avapritinib/ORY-1001 (12 nM each), or avapritinib/GSKLSD1 (12 nM each) stained with May Grunwald-Giemsa. Scale bar represents 10 μm. * $p < 0.05$, ** $p < 0.01$, *** $p < 0.001$, **** $p < 0.0001$

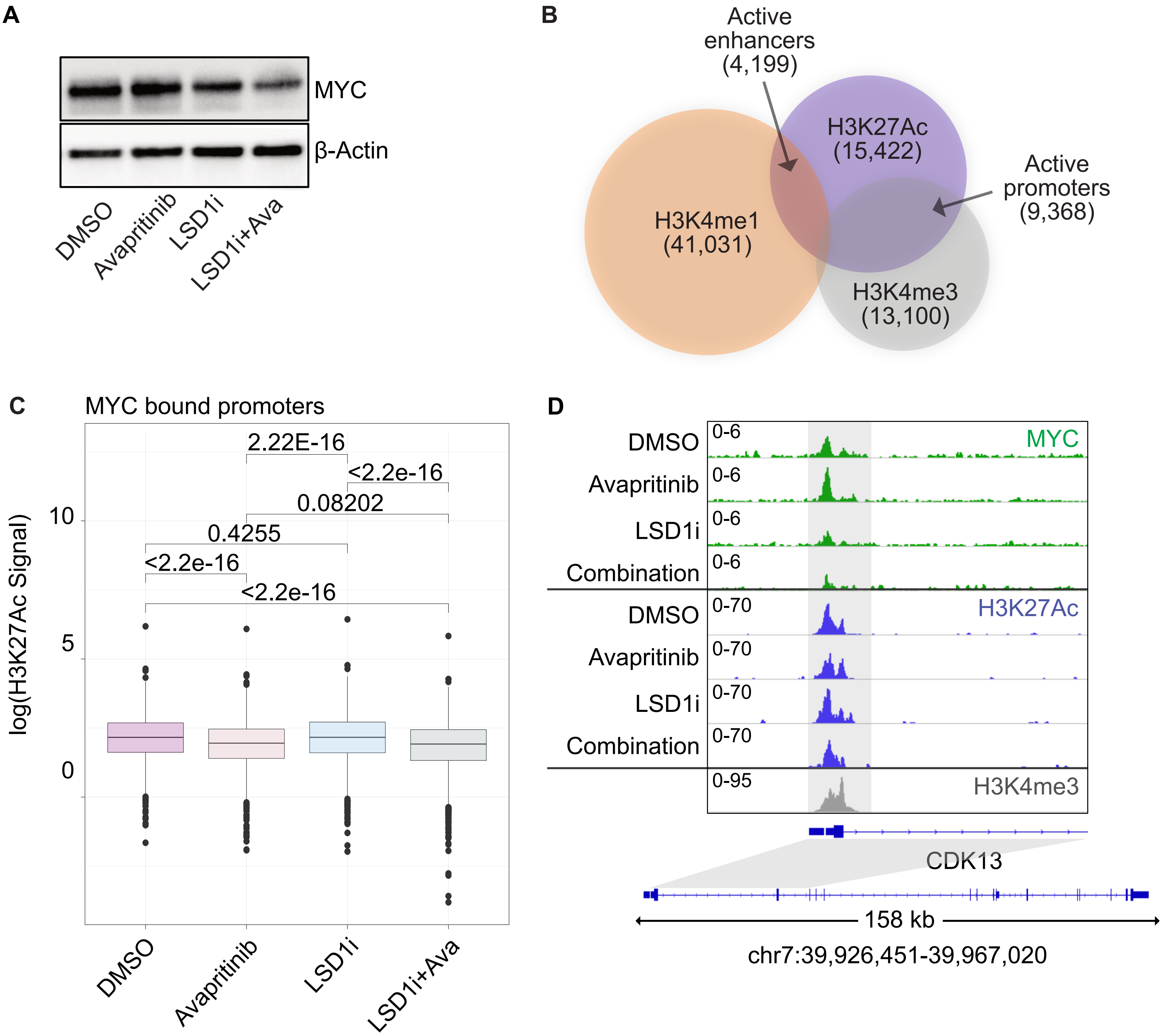


**Supplementary Figure 2. Loss of total MYC protein and acetylation at MYC bound promoters**
**A**. Western blot of total MYC and loading control (β-Actin) in Kasumi-1 cells 24 h after treatment with avapritinib (12 nM) and/or ORY-1001 (12 nM; LSD1i). **B.** Venn diagram of peaks from Kasumi-1 CUT&Tag used to define regulatory elements. Promoters defined as H3K4me3 peaks <1 kb from a TSS; enhancers defined as H3K4me1 peaks >1 kb of a TSS. Regulatory elements that overlap with H3K27Ac are defined as active (*n*=2/group). **C.** Box and whisker plot of H3K27Ac signal at MYC bound promoters. KS test used to assess statistical significance. **D.** Example of lost acetylation at MYC bound promoter of CDK13.

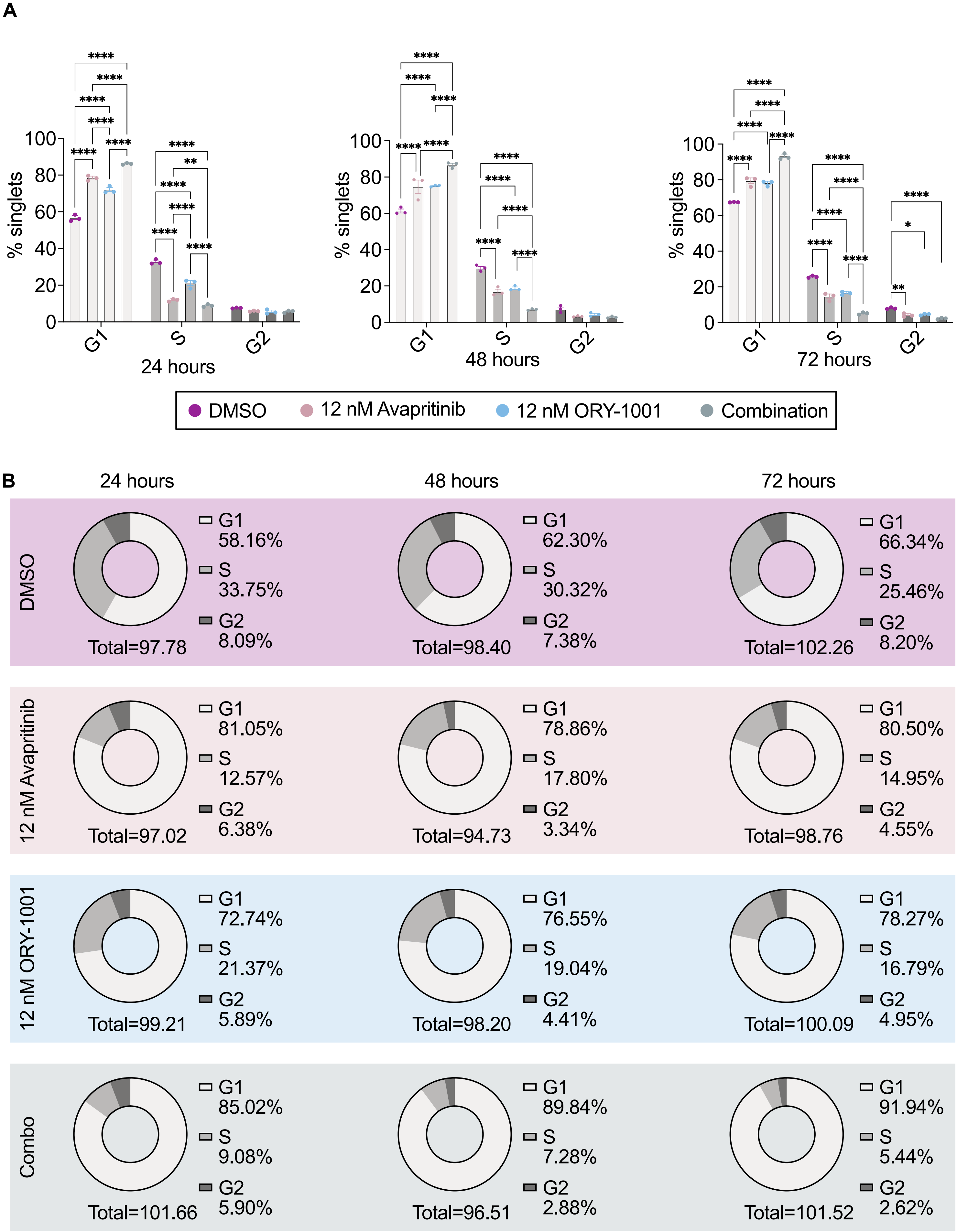
A
% singlets
100
80
60
40
20
0
****
**
*
G1
S
G2
24 hours
48 hours
72 hours
DMSO
12 nM Avapritinib
12 nM ORY-1001
Combination
B
24 hours
48 hours
72 hours
DMSO
G1 58.16%
S 33.75%
G2 8.09%
Total=97.78
G1 62.30%
S 30.32%
G2 7.38%
Total=98.40
G1 66.34%
S 25.46%
G2 8.20%
Total=102.26
12 nM Avapritinib
G1 81.05%
S 12.57%
G2 6.38%
Total=97.02
G1 78.86%
S 17.80%
G2 3.34%
Total=94.73
G1 80.50%
S 14.95%
G2 4.55%
Total=98.76
12 nM ORY-1001
G1 72.74%
S 21.37%
G2 5.89%
Total=99.21
G1 76.55%
S 19.04%
G2 4.41%
Total=98.20
G1 78.27%
S 16.79%
G2 4.95%
Total=100.09
Combo
G1 85.02%
S 9.08%
G2 5.90%
Total=101.66
G1 89.84%
S 7.28%
G2 2.88%
Total=96.51
G1 91.94%
S 5.44%
G2 2.62%
Total=101.52

**Supplementary Figure 3. Decreased cell cycle progression with combined LSD1 and KIT inhibition**

**A, B.** Cell cycle flow cytometry assay with PI of Kasumi-1 cells treated for 24, 48, or 72 h with avapritinib (12 nM) and/or ORY-1001 (12 nM). Bar chart of percentage of cells within G1, S, or G2 phase and pie chart of percent of cells within each phase of cell cycle (*n*=3/group). * $p < 0.05$, **$p < 0.01$, ****$p < 0.0001$

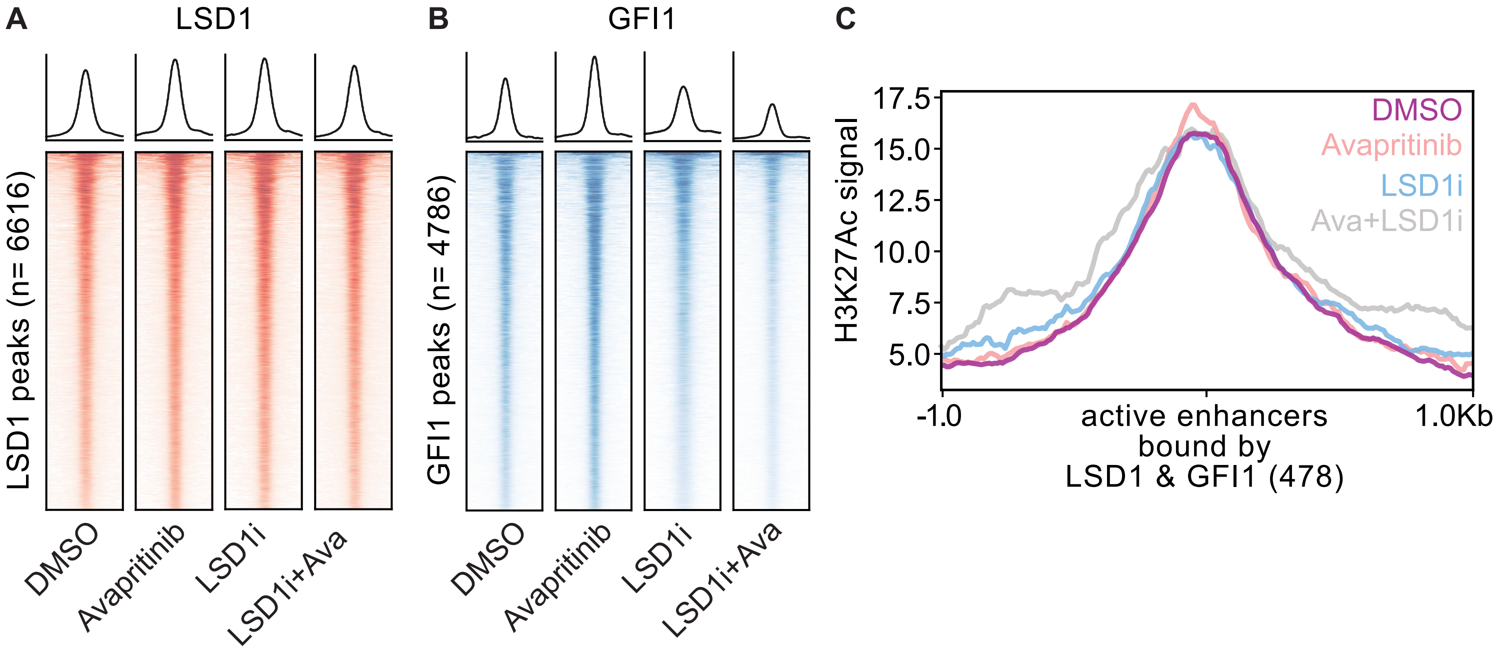


**Supplementary Figure 4. Activation of LSD1 and GFI1 bound enhancers with KIT and LSD1 inhibition**

**A, B.** Kasumi-1 cells were treated for 12 h (LSD1) or 24 h (GFI1) with avapritinib (12 nM) and/or LSD1 inhibitor (12 nM; ORY-1001 and GSK-LSD1 respectively) then subject to CUT&RUN (*n*=2-3/group). Heatmaps of global signal for LSD1 and GFI1 at their respective high confidence consensus peak sets (peak apex ± 1 kb). **C.** Peak profile and plot of H3K27Ac signal at active enhancers bound by LSD1 and GFI1 in Kasumi-1 cells after 24 hours of treatment with avapritinib (12 nM) and/or GSK-LSD1 (12 nM; LSD1i).

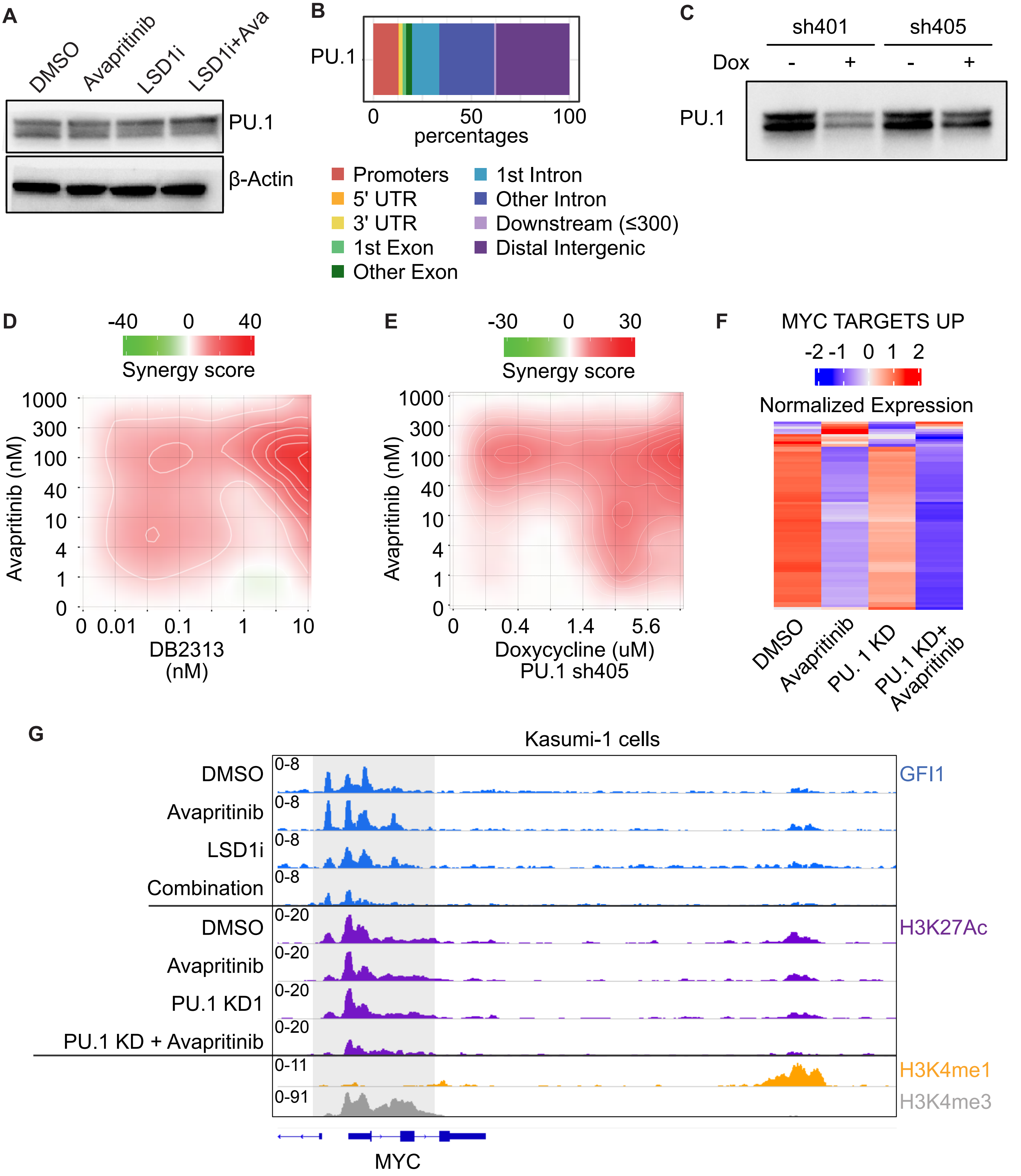


**Supplementary Figure 5. PU.1 inhibition and knock down synergize with KIT inhibition**
**A.** Western blot of total PU.1 and loading control (β-Actin) in Kasumi-1 cells following 24 h treatment with avapritinib (12 nM) and/or ORY-1001 (12 nM). **B.** Annotation of high confidence consensus PU.1 peaks. **C.** Western blot of PU.1 in Kasumi-1 shRNA knockdown cells treated with and without 1

μg/mL doxycycline for 48 h. **D, E.** Drug matrix of Kasumi-1 cells treated for 72 h with avapritinib and DB2313 or doxycycline with synergy assessed by ZIP. **F**. Normalized expression of MYC targets up gene set for DMSO, avapritinib (12 nM), PU.1 sh401 (1 μg/mL doxycycline), or both (*n*=3/group). **G.** GFI1, H3K27Ac, H3K4me1, and H3K4me3 tracks at MYC promoter and +26 kB enhancer. For GFI1 CUT&RUN, Kasumi-1 cells were treated with avapritinib (12 nM), GSK-LSD1 (LSD1i; 12 nM), or both. For H3K27Ac CUT&Tag, Kasumi-1 cells were treated with avapritinib (12 nM), PU.1 sh401 (1 μg/mL doxycycline), or both.

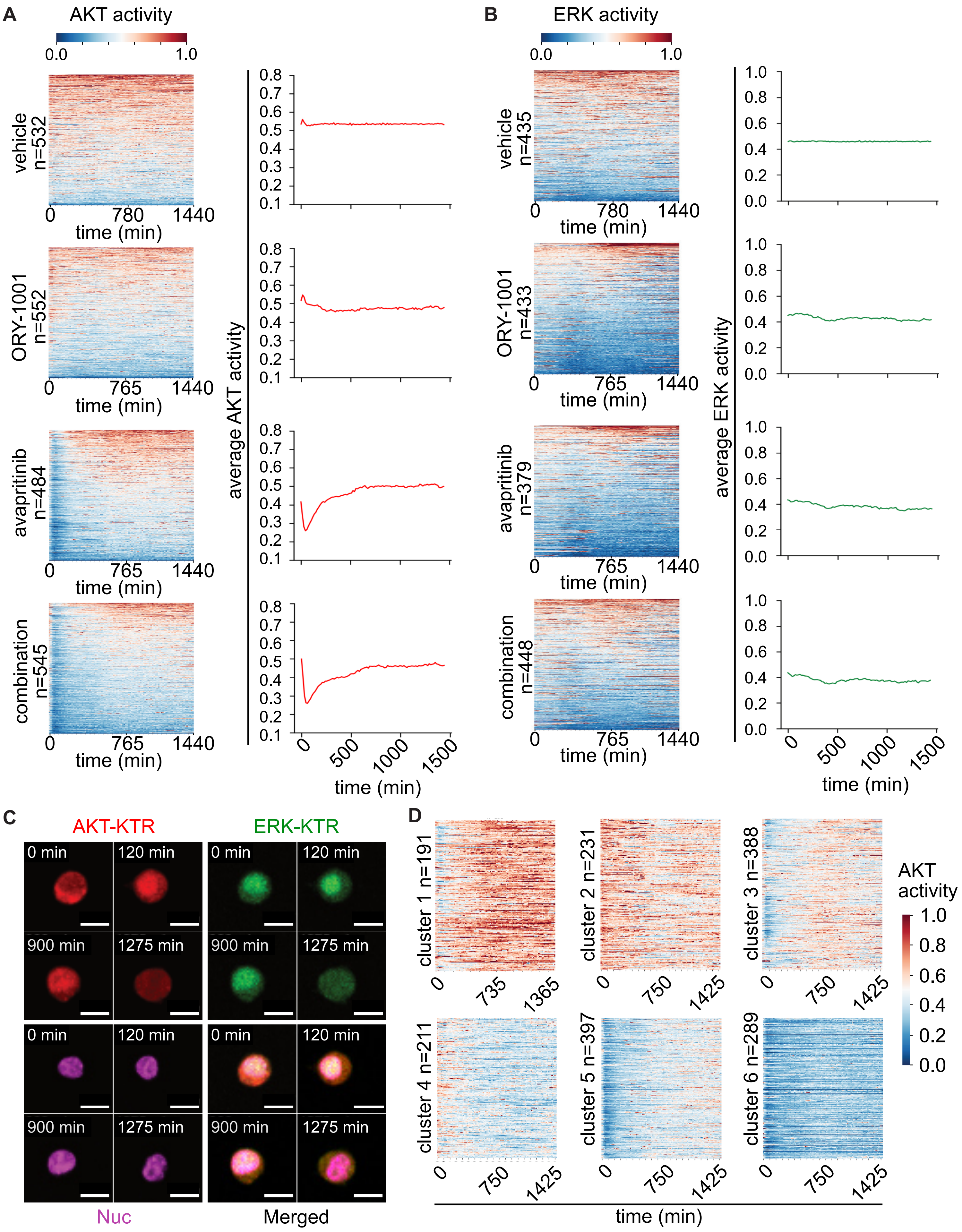
A
AKT activity
0.0
1.0
vehicle n=532
ORY-1001 n=552
avapritinib n=484
combination n=545
0
780
765
1440
time (min)
average AKT activity
0.8
0.7
0.6
0.5
0.4
0.3
0.2
0.1
0
500
1000
1500
B
ERK activity
vehicle n=435
ORY-1001 n=433
avapritinib n=379
combination n=448
average ERK activity
1.0
0.8
0.6
0.4
0.2
0.0
C
AKT-KTR
ERK-KTR
0 min
120 min
900 min
1275 min
Nuc
Merged
D
cluster 1 n=191
cluster 2 n=231
cluster 3 n=388
cluster 4 n=211
cluster 5 n=397
cluster 6 n=289
735
1365
750
1425
AKT activity

**Supplementary Figure 6. Dual LSD1 and KIT inhibition leads to reduction of AKT and not ERK signaling**

**A, B.** Heatmaps of AKT and ERK dynamics of individual cells (every row) in response to ~25h treatment of DMSO, ORY-1001 (12 nM), avapritinib (100 nM), and the combination. *n* represents the total number of tracked cells in each condition. Lineplots represent population average of AKT and ERK dynamics over time. **C.** Example of time-series images of fluorescent cells under the combination treatment. AKT-KTR = mScarlet, ERK-KTR = Clover, and Nuc = miRFP670nano. Scale bar represents 10 μm. **D.** Heatmaps of cells grouped into different AKT dynamic clusters by hierarchical clustering.

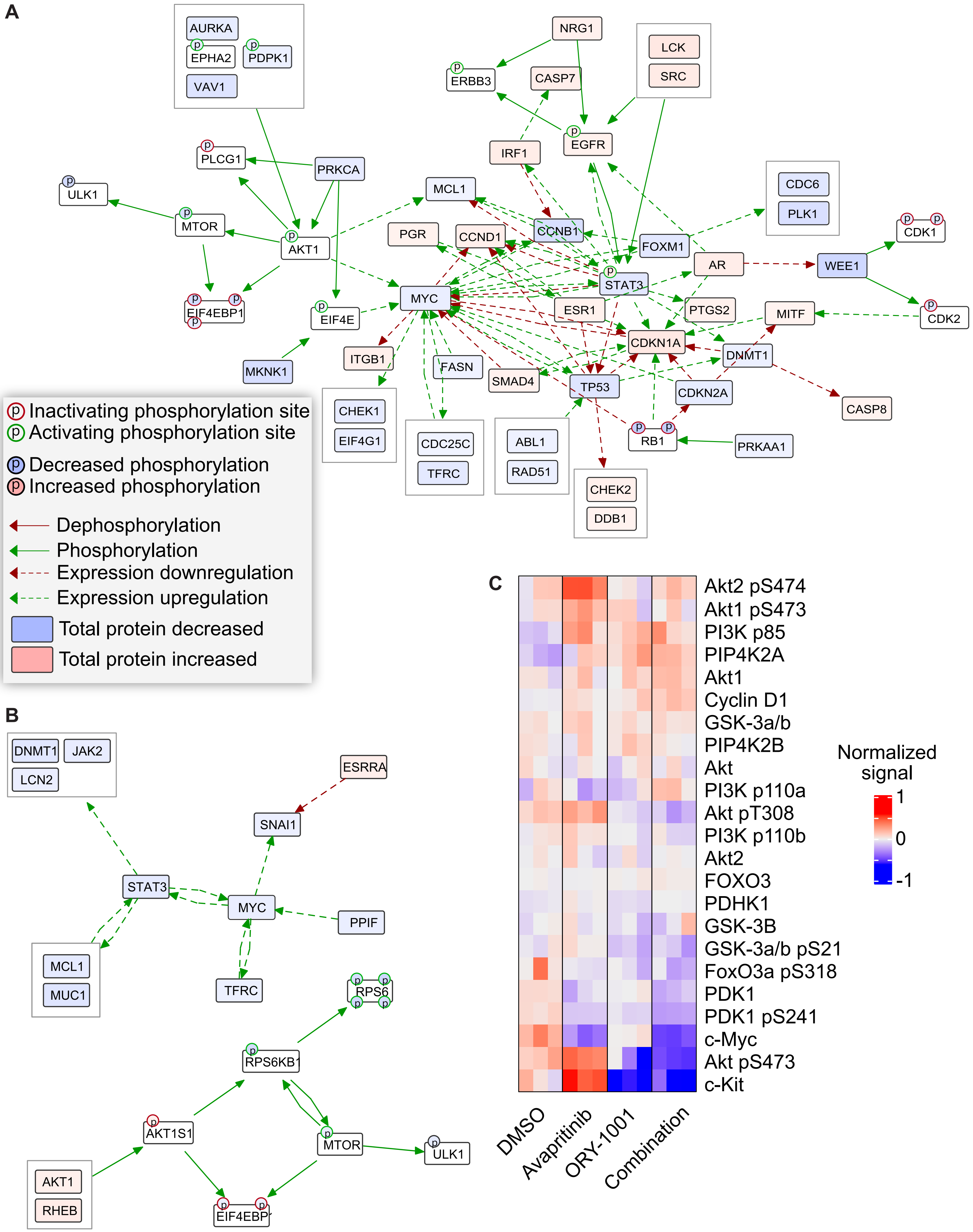
A
AURKA
EPHA2
PDPK1
VAV1
NRG1
ERBB3
CASP7
LCK
SRC
PLCG1
PRKCA
ULK1
MTOR
AKT1
EIF4EBP1
EIF4E
MKNK1
IRF1
EGFR
MCL1
CCNB1
PGR
CCND1
FOXM1
AR
WEE1
CDC6
PLK1
CDK1
CDK2
MYC
STAT3
ESR1
PTGS2
MITF
CDKN1A
DNMT1
ITGB1
FASN
SMAD4
TP53
CDKN2A
CASP8
RB1
PRKAA1
CHEK1
EIF4G1
CDC25C
TFRC
ABL1
RAD51
CHEK2
DDB1
Inactivating phosphorylation site
Activating phosphorylation site
Decreased phosphorylation
Increased phosphorylation
Dephosphorylation
Phosphorylation
Expression downregulation
Expression upregulation
Total protein decreased
Total protein increased
B
DNMT1
JAK2
LCN2
ESRRA
SNAI1
STAT3
MYC
PPIF
MCL1
MUC1
TFRC
RPS6
RPS6KB1
AKT1S1
MTOR
ULK1
AKT1
RHEB
EIF4EBP
C
Akt2 pS474
Akt1 pS473
PI3K p85
PIP4K2A
Akt1
Cyclin D1
GSK-3a/b
PIP4K2B
Akt
PI3K p110a
Akt pT308
PI3K p110b
Akt2
FOXO3
PDHK1
GSK-3B
GSK-3a/b pS21
FoxO3a pS318
PDK1
PDK1 pS241
c-Myc
Akt pS473
c-Kit
Normalized signal
1
0
-1
DMSO
Avapritinib
ORY-1001
Combination

**Supplementary Figure 7. Identify AKT as prominent pathway leading to MYC repression**
**A**. CausalPath analysis of RPPA results of Kasumi-1 cells treated for 24 h with avapritinib (100 nM) and/or ORY-1001 (12 nM) or DMSO. **B.** CausalPath analysis of RPPA as in A after 1 h of drug exposure. **C**. Heatmap of the normalized signal for PI3K/AKT pathway members after 24 h of drug exposure.

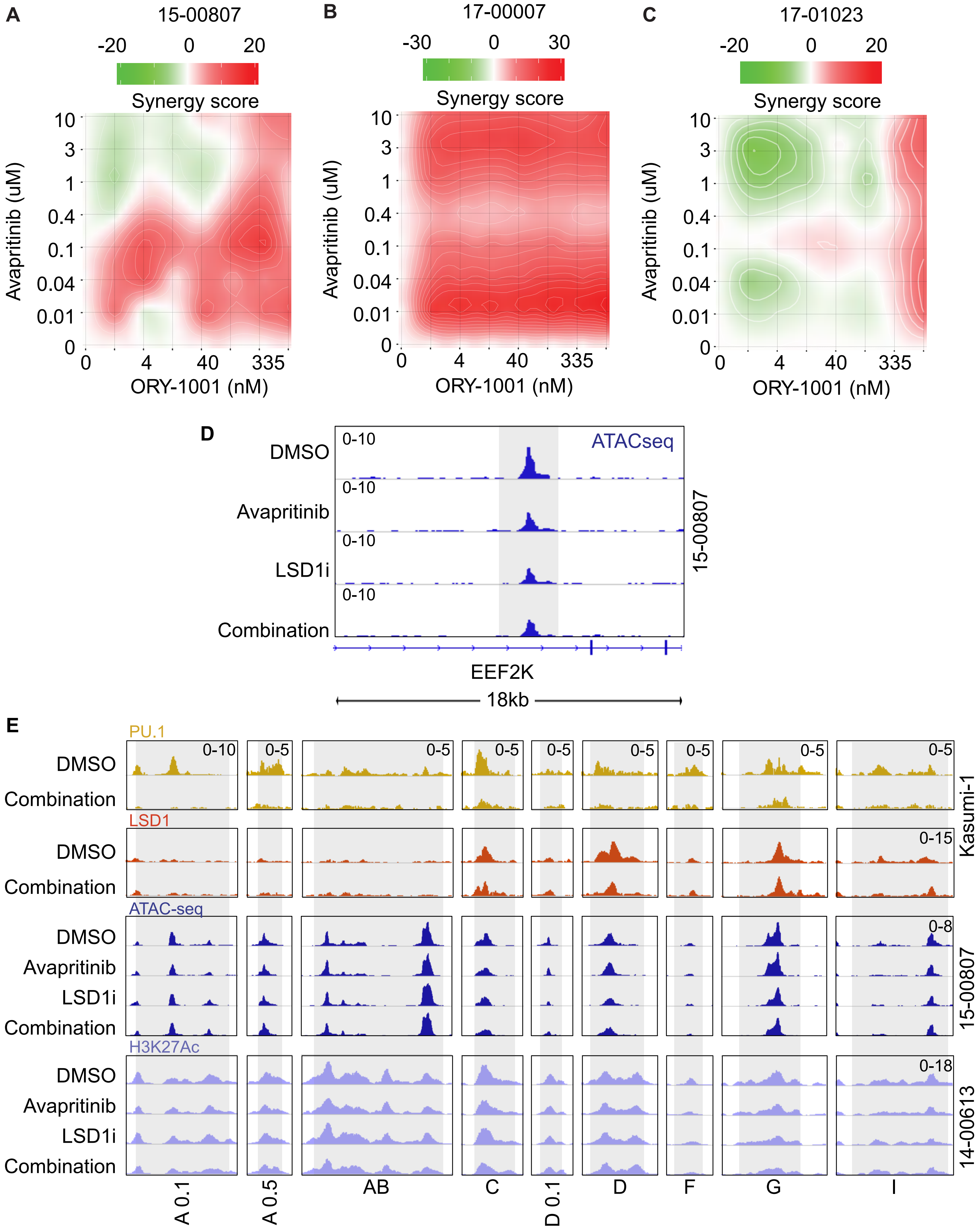
A
15-00807
-20 0 20
Synergy score
Avapritinib (uM)
ORY-1001 (nM)
B
17-00007
-30 0 30
Synergy score
Avapritinib (uM)
ORY-1001 (nM)
C
17-01023
-20 0 20
Synergy score
Avapritinib (uM)
ORY-1001 (nM)
D
ATACseq
DMSO
Avapritinib
LSD1i
Combination
0-10
15-00807
EEF2K
18kb
E
PU.1
LSD1
ATAC-seq
H3K27Ac
DMSO
Avapritinib
LSD1i
Combination
Kasumi-1
15-00807
14-00613
A 0.1
A 0.5
AB
C
D 0.1
D
F
G
I

**Supplementary Figure 8. Additional KIT and LSD1 inhibitor synergy plots in KIT-mutant AML patient samples**

**A-C.** Drug matrices of patient samples 15-00807, 17-00007, and 17-0123 treated for 72 h with avapritinib and ORY-1001 with synergy assessed by ZIP score. **D.** Example of decreased accessibility at cell cycle associated locus, EEF2K, from bulk ATAC-seq in 15-00807. **E.** Visualization of Kasumi-1 PU.1 and LSD1, 15-00807 bulk ATAC-seq, and 14-00613 H3K27Ac at entire BENC locus.

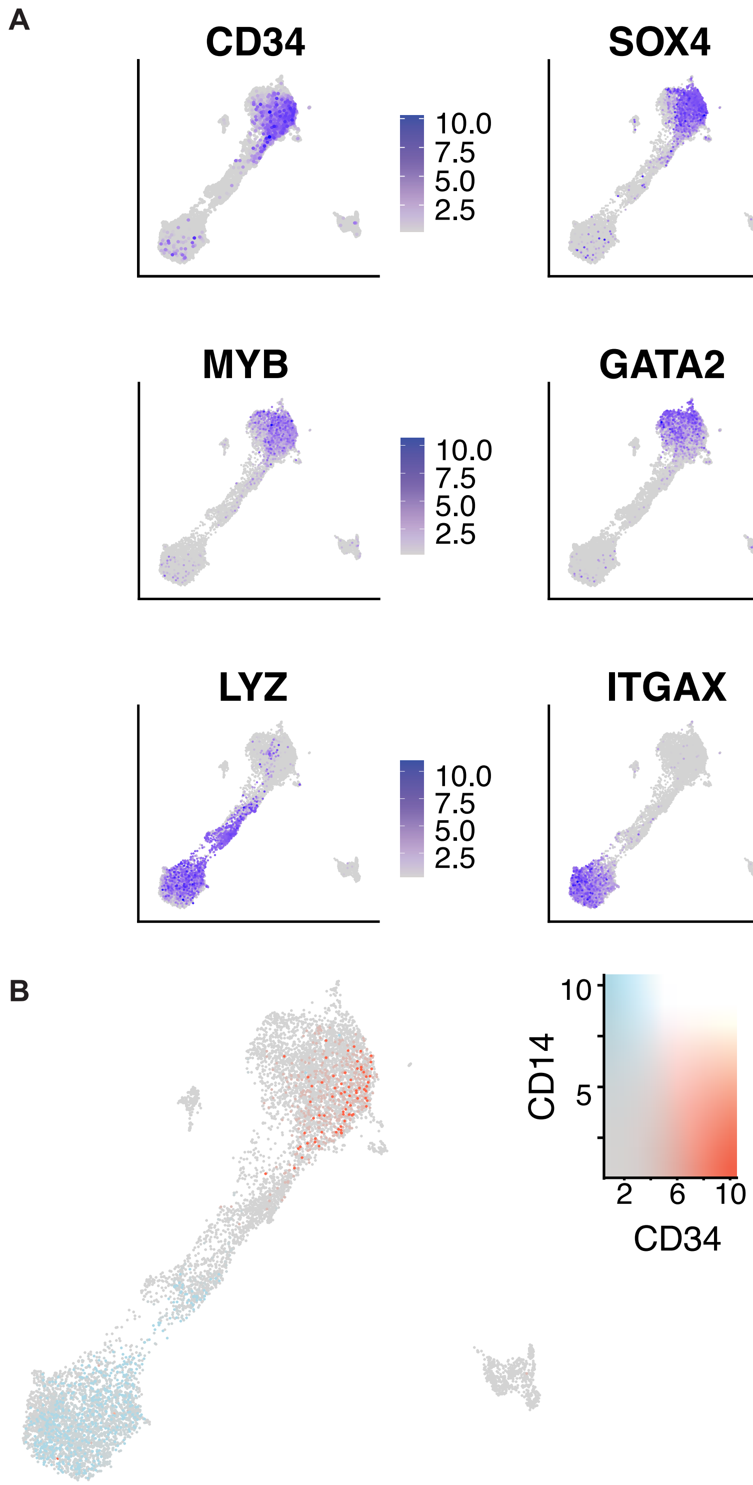


**Supplementary Figure 9. Single cell RNA-seq markers for differentiation**
**A.** Integrated single cell RNA-seq object showing the expression of differentiation markers. Markers were used to identify early, mid, and late stages of hematopoiesis in patient sample 14-00613. **B.** Detected levels of CD34 (early) and CD14 (late) in integrated object to show overall differentiation trajectory.
